# Identifying control strategies to eliminate African swine fever from the United States swine industry in under 12 months

**DOI:** 10.1101/2024.12.17.628884

**Authors:** Abagael L. Sykes, Jason A. Galvis, Kathleen C. O’Hara, Lindsey Holmstrom, Cesar Corzo, Gustavo Machado

## Abstract

With the rising risk of African swine fever (ASF) introduction into the U.S., there is substantial emphasis on preparation for an epidemic to mitigate the economic and societal impacts. Mathematical models represent a vital tool for simulating future epidemics and examining the effectiveness of response strategies. This study expands on our spatially explicit stochastic compartmental farm-level transmission model, *PigSpread-ASF*, to identify the control strategies necessary to eliminate ASF from domestic swine populations in three, six, nine, and twelve months. We achieved this by incrementally increasing the intensity of the control actions detailed in the USDA’s national response plan, which consists of i) quarantine and depopulation of detected farms; ii) a 72-hour movement standstill; iii) contact tracing with subsequent diagnostic testing; and iv) the implementation of control areas (infected and buffer zone) and surveillance zones (including routine diagnostic testing, pre-permit testing and movement restrictions). For this model, ASF was deemed eliminated after three consecutive months of no new ASF cases following each time period, as determined by WOAH.

Our results indicate that the current national response plan would need to i) increase radii and duration of control areas and surveillance zones, ii) extend the traceback and quarantine for contact farms; iii) extend the movement standstill; iv) prohibit repopulation of depopulated farms; and v) quicker baseline detection of ASF, at varying intensities, to eliminate ASF within three, six, nine and twelve months. The elimination of ASF in 12-months required extension of the buffer zone radius to 5 km and maintenance of the control areas and surveillance zones for 60 days, a traceback and quarantine of 60 and 30 days for the contact tracing, and a standstill of 30 days. In contrast, the three-month scenario required extension of the infected zone, buffer zone and surveillance zone radii to 20 km each and maintenance of the control area and surveillance zones for 90 days, a traceback and quarantine of 90 days for contact tracing, and a standstill of 90 days. By intensifying the current national response plan, ASF would likely be eliminated within 12-months of its introduction. However, it is pertinent to consider the limitations posed by resource capacities and the impact that intensifying control may have on business continuity. Nevertheless, our study provides beneficial guidance to aid preparation for a future ASF introduction and estimates the infrastructure and personnel required to bring an epidemic under control promptly.

## 1. Introduction

The risk of African swine fever (ASF) being introduced into the U.S. through both legal and illegal importation of infected pork products (Fanelli et al., 2022; Herrera-Ibatá et al., 2017; Jurado et al., 2019; Schambow et al., 2022b) is significant given the continued outbreaks throughout Asia, Europe and the island of Hispaniola (FAO, 2023; Gonzales et al., 2021; Schambow et al., 2022a; Schettino et al., 2023). An ASF outbreak could have harsh economic consequences and disrupt the U.S. swine industry (Carriquiry et al., 2020; Sykes et al., 2023). With no approved, effective, commercially available vaccine (Chathuranga and Lee, 2023; Penrith and Kivaria, 2022; Urbano and Ferreira, 2022), mitigation of a future epidemic relies primarily on improved knowledge of ASF infection dynamics within the U.S. and assessment of the available control and eradication actions (Hayes et al., 2021; Penrith and Kivaria, 2022).

In our previous work, we developed a stochastic mathematical ASF farm-level transmission model (Sykes et al., 2023) to investigate the trajectory of a U.S. ASF epidemic and assess the effectiveness of five control and eradication actions based on the control strategies detailed in the United States Department of Agriculture (USDA) national ASF response plan, “The Red Book” (USDA, 2023a). The control actions included i) quarantine and depopulation of detected farms; ii) a 72-hour movement standstill for all movements of pigs and vehicles; iii) contact tracing of direct (pig movements) and indirect contacts (vehicle movements); and iv) the implementation of control areas (infected and buffer zones) and surveillance zones. Though not included in the Red Book the efficacy of depopulation of high-risk farms (i.e., direct contacts) was also assessed (Sykes et al., 2023). Our model provided estimates of the expected spread dynamics and the effectiveness of the current control actions, and also indicated that elimination of ASF may take longer than four months, with only 29% of simulations reporting elimination by day 140 (Sykes et al., 2023). To mitigate the financial and societal impacts of ASF (Alarcon et al., 2014; Carriquiry et al., 2020; Cooper et al., 2022; Dixon et al., 2020), an introduction into the U.S. must be detected, controlled and eradicated swiftly. Therefore, it is vital to identify the strategies necessary to eliminate ASF before the virus becomes widespread within the U.S. commercial swine population, while examining the feasibility and availability of required resources (e.g., the volume of samples, permits and depopulated animals).

In this study, we revised our stochastic farm-level mathematical model (*PigSpread-ASF)* (Sykes et al., 2023); we updated the control actions to match the 2023 national response plan (USDA, 2023a) and intensified them to eliminate an ASF epidemic in the U.S. within three, six, nine and twelve months after virus introduction. The control actions investigated were i) quarantine and depopulation of detected farms; ii) 72-hour movement standstill; iii) contact tracing; and iv) implementation of control areas and surveillance zones. The present work extends our current ASF model (Sykes et al., 2023) by aligning it to the national response plan (USDA, 2023a), thus increasing its potential applications for decision-making in the event of an ASF introduction.

## 2. Materials and Methods

### 2.1 Population data

Farm population and geolocation data, including the production type and capacity, were provided for 1,981 commercial farms from eight swine production companies across three U.S. states. Farms were identified by their unique national premises identification number. When multiple farms were registered under the same identification number, they were considered one farm. Farms were grouped into eight production types based on the production stage of pigs present at the farm: sow, nursery, finisher, gilt development, boar stud, wean-to-finish, farrow-to-finish, and isolation. The farm’s capacity data was used to calculate depopulation and diagnostic testing requirements under the relevant control scenarios.

### 2.2 Model structure

Briefly, the *PigSpread-ASF* model is a stochastic, farm-level, compartmental transmission model with four health states: Susceptible (S), Exposed (E), Infected (I), and Detected (D) (Sykes et al., 2023). Transmission routes included i) movement of pigs between farms; ii) local spread, reflecting transmission related to spatial proximity; iii) vehicles moving pigs between farms (pig trucks); iv) vehicles moving pigs from farms to slaughterhouses (market trucks); v) vehicles delivering feed to farms (feed trucks); and vi) vehicle movements between farms without a defined role (undefined trucks) (Galvis and Machado, 2023). The model begins with one initial farm infected, randomly selected from the study population. The probability of transition from S to E compartment (Figure 1), *Y_it_*, is defined by the force of infection from vehicle movements, local spread, and movements of exposed pigs (Supplementary Material Equations 1 to 14). Movements of pigs from an infected or detected farm contributed to the probability of farms transitioning into the I and/or D compartments, represented by *V_it_* and *W_it_*, respectively. Farms also transition from compartment E to compartment I through the development of infectiousness 1/σ, where σ is the latent period of the ASF virus. Farms also transition from compartment I to compartment D via detection based on above normal mortality, determined by the baseline detection rate,

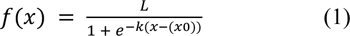

where *L* represented the effectiveness of surveillance for the production type; *k*, the logistic growth rate; *x*, the time spent in the I compartment; and *x*0, the time taken to reach the midpoint of the logistic curve (equivalent to half of the average time to detection). Additional details of the model structure are described in-depth in our previous publication (Sykes et al., 2023).

**Figure 1.**
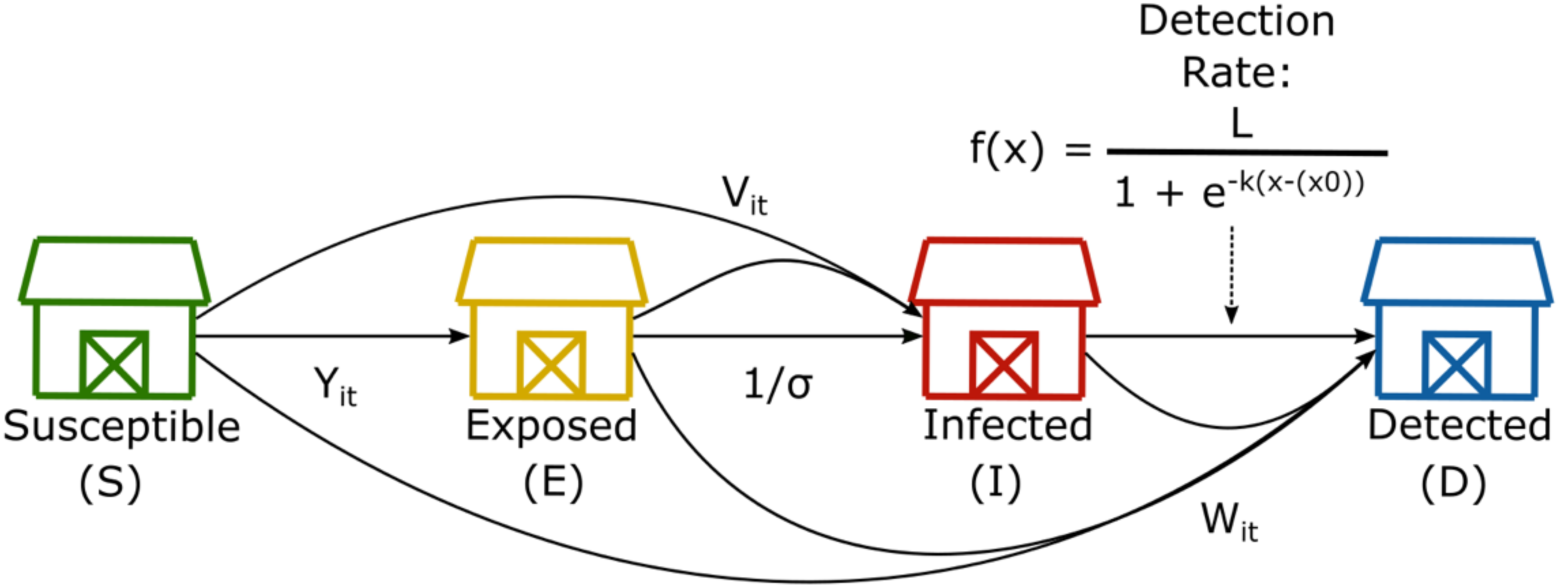
Schematic representation of the transitions between the different health compartments of our SEID transmission model. *Y*_it_, *V*_it_, and *W*_it_ represent the probability of moving into the E, I, or D compartments, respectively, through receipt of an exposed, infected, or detected movement of swine. σ represents the latent period of ASF virus. Within the baseline detection rate, L indicates the surveillance effectiveness (determined by production type), *k* represents the logistic growth rate, and *x*0 indicates the average time in days to reach the midpoint of the logistic curve (equivalent to half of the average time to detection) (Supplementary Material Equations 1 to 14).

#### Animal movement networks

As described in our previous publication (Sykes et al., 2023), pig movements are represented by directed, unweighted temporal networks of swine movements. If more than one shipment of pigs occurred between the same origin and destination farm on a single day, they were collated and considered one movement.

#### Vehicle movement networks

In this revised model, the directed vehicle networks for the movements of pig trucks, market trucks, feed trucks, and undefined trucks were reconstructed as described elsewhere (Galvis and Machado, 2023). Briefly, the GPS locations of the vehicles and the outline of the farm’s perimeter buffer area (PBA) (Supplementary Material Figure S1), provided in their Secure Pork Supply plans, were used to determine the visits a vehicle made. A vehicle had to be within 50 m of a farm’s PBA while registering a speed of zero km/h for five minutes to be considered as “visiting” a farm (Galvis and Machado, 2023). If PBA information was not available, the coordinates associated with the farms were used instead. We considered vehicles as contaminated after visiting a simulated ASF-positive farm. The force of infection for the movement of the vehicle was weighted according to pathogen stability (Galvis and Machado, 2023), calculated as a function of the vehicle travel time between the two farms or nodes, and the average environmental temperature during the journey (Galvis and Machado, 2023). A stability of one indicated high pathogen stability, while a stability of zero indicated low pathogen stability. Additionally, we included vehicle movements to cleaning stations, defined as being within 500 m of a cleaning station for at least 60 minutes. Cleaning effectiveness for these visits was set at 50% and was compared to a stochastic threshold, above which the pathogen was assumed to have been eliminated from the vehicle. In such cases, these vehicles had no onward transmission (Galvis and Machado, 2023).

#### Local spread

The dynamics of local spread are included in the model to account for transmission through routes that cannot be captured explicitly, including movements of personnel, movements of equipment and movements of wildlife such as feral swine (Bellini et al., 2016; Chuchard et al., 2022; Gao et al., 2021; Miller et al., 2017; Podgórski and Śmietanka, 2018; Yoo et al., 2021; Zani et al., 2019). In this model interaction we updated local spread from a gravity model to a transmission kernel in which the probability of infection decreases with increased distance (Equation 2 and Supplementary Material Figure S2) (Cardenas et al., 2022).

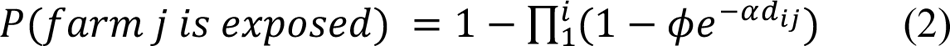

Here, farm *i* represents an infected and/or detected farm, and farm *j* represents a susceptible farm. The probability of farm *j* becoming exposed was calculated from the inverse probability of farm *j* not becoming exposed by infected farms nearby. We implemented a distance cut-off from 1 km to 10 km for the local spread to reflect the uncertainty about ASF local dissemination dynamics (Andraud et al., 2019; Boklund et al., 2020; Bradhurst et al., 2021; Chenais et al., 2019; Fodor et al., 2015; Halasa et al., 2016; Iglesias et al., 2016; Pepin et al., 2021). The transmission kernel (Equation 2) was shaped by parameters Φ and α that control the maximum probability of transmission and the steepness with which the probability declines with increasing distance (Cardenas et al., 2022). The parameters Φ and α values were calibrated using historic porcine epidemic diarrhea virus (PEDV) data as a proxy for ASF, described elsewhere (Sykes et al., 2023). In the model, it was assumed that if a farm was ASF positive (either infected or detected) it was capable of local spread to farms within the cutoff distance.

### 2.3 Model calibration

Our parameters were calibrated using historic PEDV outbreak data and an Approximate Bayesian Computation (ABC) rejection algorithm (Minter and Retkute, 2019), as implemented in our previous model (Sykes et al., 2023). The PEDV historical data was taken from the beginning of the 2013-2014 PEDV epidemic. In 2013, PEDV was a novel virus spreading in a susceptible U.S. swine population (Stevenson et al., 2013), providing an appropriate proxy for ASF introduction.

Using the ABC algorithm, we identified values of the transmission rate parameters (β_n_, β_s_, β_p_, β_m_, β_f_, and β_b_), the local spread kernel parameters (α and Φ), the kernel cut-off distance, and the effectiveness of passive surveillance (*L*_2_, where *z* indicates the production type) that replicated the temporal distribution of the observed PEDV cases (Table 1). The prior distributions applied to the ABC algorithm are presented in the Supplementary Material, Table S1. From the accepted ABC simulations we extracted the sets of parameter values provided, hereafter referred to as particles, which were subsequently used in our model as the calibrated parameter values.

**Table 1.** Summary of parameter values extracted from accepted simulations via an Approximate Bayesian Computation (ABC) rejection algorithm.

| Parameter | Notation | Median particle value (IQR) |
| --- | --- | --- |
| Transmission rate for infected pig movements | $\beta_s$ | 1.51 (1.50 - 1.52) |
| Transmission rate for exposed pig movements | $\beta_n$ | 1.51 (1.49 - 1.52) |
| Transmission rate for movements of pig trucks | $\beta_p$ | 0.0614 (0.0594 - 0.0627) |
| Transmission rate for movements of market trucks | $\beta_m$ | 0.0606 (0.0576 - 0.0628) |
| Transmission rate for feed delivery trucks | $\beta_f$ | 0.0288 (0.0283 - 0.0293) |
| Transmission rate for undefined trucks | $\beta_b$ | 0.0605 (0.0589 - 0.0621) |
| Maximum probability of transmission | $\phi$ | 0.0630 (0.0620 - 0.0643) |
| The gradient of transmission probability decay | $\alpha$ | 1.39 (1.39 - 1.39) |
| Effective surveillance in sow farms | $L_{sow}$ | 0.851 (0.769 - 0.929) |
| Effective surveillance in nursery farms | $L_{nursery}$ | 0.634 (0.586 - 0.689) |
| Effective surveillance in finisher farms | $L_{finisher}$ | 0.556 (0.544 - 0.576) |

Values have been rounded to three significant figures.

### 2.4 Control actions

In our previous *PigSpread-ASF* model iteration (Sykes et al., 2023), we implemented the following five control and eradication actions: i) quarantine and depopulation of detected farms; ii) a 72-hour movement standstill restricting all movements; iii) contact tracing of direct contacts (via animal movements) and indirect contacts (via vehicle movements); iv) depopulation of direct contacts; and v) implementation of control areas and surveillance zones (USDA, 2023a). In the revised model, we made the following changes:

#### Quarantine and depopulation of detected farms

As described in our previous model iteration, detected farms are quarantined and scheduled for depopulation (Sykes et al., 2023). The delay between detection and depopulation is based on the number of farms awaiting depopulation and the depopulation limit of four farms per day. Once depopulation is complete, farms remain empty for 30 days to allow for cleaning and disinfection before being repopulated, which is twice the time required for urine and feces to cease being infectious at 4°C (Davies et al., 2017).

#### Movement standstill of live pigs and associated vehicles

The 72-hour movement standstill was restricted to movements of live swine only, which included animal movements, pig trucks, market trucks, and undefined trucks (USDA, 2023a). This is implemented upon the initial detection of an ASF-positive farm in the study area and is applied to all farms. Standstill compliance was assumed to be 100%.

#### Contact tracing

Contacts resulting from swine movements (direct contacts) and vehicle movements (indirect contacts) to and from detected farms within the previous 30 and 15 days, respectively, were quarantined for 15 days starting the following day, stopping all movements on and off the farm (USDA, 2023a). Efficacy of contact tracing was assumed to be 100%. During this quarantine period, farms were required to test for ASF at regular intervals (see Supplementary Material, Table S2). Diagnostic testing increases the chance of detection in addition to the already present baseline detection rate.

#### Depopulation of high-risk contacts (direct contacts)

Only farms that tested positive for ASF were depopulated, while direct contact farms were quarantined but not depopulated (USDA-APHIS CEAH, 2023).

#### Control areas and surveillance zones

Farms within the control area, consisting of a 3 km infected zone and a 2 km buffer zone surrounding infected farms, were quarantined immediately. The quarantine was lifted from each farm once they could present one negative ASF test result, however, farms were still required to be tested at regular intervals (see Supplementary Material, Table S2) until the control area and surveillance zone were lifted after 30 days. The daily processing capacity for blood samples (pooled by five) was limited to 25 farms for North Carolina, ten farms for Virginia, and ten farms for South Carolina based on current laboratory capacities (State Animal Health Official, personal correspondence, 2023).

Additionally, farms were required to test negative for ASF three days and one day before moving pigs, as described in USDA’s published permitting templates and hereafter referred to as pre-permit testing. Pigs were prohibited from moving if farms did not meet this requirement or tested ASF positive; the regular surveillance testing required as part of the control area and surveillance zones did not contribute to the pre-movement testing requirements. Per barn, 31 samples were required and could be pooled into groups of five; for each movement, we assumed that all barns were being moved. Individual one-movement permits continued to be required for swine movements and movements of market trucks, while permits for swine trucks were removed as they were covered by the swine movement permit. For the movements of feed trucks, 30-day permits were allowed for recurring movements between farms (USDA, personal correspondence, 2023).

As in our previous model, control actions were triggered after detection of the first case of ASF in the study population. We assumed 100% public compliance with the control actions (Sykes et al., 2023) and the sensitivity of diagnostic testing was fixed at 95% (Pikalo et al., 2022).

### 2.5 ASF elimination scenarios

We investigated alternative control strategies to eliminate ASF from the study population in three, six, nine, and twelve months^1^. We incrementally intensified our control actions to maintain business continuity, which is defined as a farm’s ability to receive feed and move swine to maintain production. The elimination control scenarios implemented at least one of the following increased controls: i) reducing the time to baseline detection for ASF; ii) increasing the capacity limits for depopulation and diagnostic testing; iii) increasing the duration of the national movement standstill; iv) increasing the timelines for traceback from and quarantine of contact farms; v) expanding the radius and duration of implementation of control areas and surveillance zones; vi) increasing the number of tests required for movements to resume within the control area; and vii) increasing the duration a depopulated farm must remain empty before repopulation. ASF elimination was determined by three consecutive months (90 days) without new ASF cases (WOAH, 2022). We defined an effective elimination strategy as eliminating ASF from 99% of simulations to account for potential outlier simulations. We simulated the national response plan control actions (USDA, 2023a), hereafter referred to as the national response plan (NRP) scenario, to compare our alternative elimination scenarios.

### 2.6 Outputs

The primary output of this study is the number of infections at 90, 180, 270, and 360 days for the elimination of ASF at three, six, nine, and twelve months after the introduction of ASF. In addition, we also included the following outputs: i) the number of diagnostic tests required (Supplementary Material Equation 15); ii) the number of permits and pre-permit tests required; and iii) the number of pigs depopulated, which were calculated using for the elimination time-frame and the additional 90 days of no ASF cases required by the end of the simulation (i.e. 180, 270, 360 and 450 days for the three, six, nine and twelve months respectively). We account for the additional 90 days for the control resource outputs because some controls will remain in place even if cases are no longer appearing.

### 2.7 Sensitivity analysis

Here, we examined the sensitivity of the secondary cases of ASF by varying the values of the following parameters: i) transmission rates (β_n_, β_s_, β_p_, β_m_, β_f_, and β_b_) ii) spatial kernel characteristics (α and Φ); iii) virus latent period (σ); iv) average time to reach the midpoint of the logistic curve (*x*0) (equivalent to half the time to detection); and v) surveillance effectiveness (*L*_2_, where *z* indicates the production type). This was achieved through Latin Hypercube sampling and partial rank correlation coefficient analysis using the R packages *pse* (Chalom et al., 2013) and *epiR* (Stevenson, 2024), with 5000 sets of parameter values. The input parameter probability density functions are displayed in Supplementary Material Table S3. Additionally, limitations related to the use of LHS-PRCC to analyze sensitivity are presented in the Supplementary Material.

## 3. Results

### 3.1 Descriptive analysis

Of the 1,981 farms included in our model, 54.4% were finishers, 19.6% were nurseries, 14.9% were sow farms, 9.0% were wean-to-finishers, 0.8% were boar studs, 0.8% were gilt development sites, 0.4% were farrow-to-finishers and <0.1% were isolation units. The majority of farms were located in North Carolina (95.8%), followed by Virginia (2.5%) and South Carolina (1.7%). The median pig capacity of farms was 3,672 (interquartile range (IQR): 2,500 - 5,760), and the median distance between farms was 69.2 km (IQR: 40.4 km - 119.3 km).

Three companies provided animal movement data for the 2020 calendar year, which equated to 29,169 animal movements with a daily median of 91 (IQR: 11.25 - 125.00) movements. The majority of movements (61.6%) originated from sow farms, followed by nursery farms (29.2%), finisher farms (6.0%), gilt development farms (2.0%), wean-to-finish farms (1.0%), and farrow-to-finish farms (0.7%). Nursery farms and finisher farms were the most common movement destinations, accounting for 40.6% and 30.8% of movements, respectively, followed by sow farms (18.1%), wean-to-finish farms (9.3%), gilt development farms (0.7%), boar-stud farms (0.4%) and farrow-to-finish farms (0.1%).

Additionally, data comprising 69,017 vehicle movements were provided by two companies, where the daily median movement was 157.5 (IQR: 74.00 - 296.25). Feed trucks accounted for 63.9% of the total movements, followed by pig trucks, which accounted for 20.5%; undefined trucks, which accounted for 13.7%; and market trucks, which accounted for 1.9%. The median number of movements per day for each vehicle movement type was 107.5 (IQR: 51.0 - 182.5) for feed trucks, 26 (IQR: 2.00 - 69.75) for pig trucks, 23 (IQR: 4.25 - 42.00) for undefined trucks, and 2 (IQR: 1 - 6) for market trucks.

### 3.2 Calibration of parameter values

As a result of the ABC rejection algorithm, we collected 98 accepted particles from 396,200 simulations (0.02% acceptance rate) (Table 1).

### 3.3 National Response Plan (NRP) scenario

The results of the NRP scenario show a median of 7 (IQR: 1 - 48) infections (including the initial seed farm) at day 90, 7 (IQR: 1 - 237) at day 180, 7 (IQR: 1 - 581) at day 270, and 7 (IQR: 1 - 1,002) at day 360. At day 360, finisher farms accounted for the most infections with a cumulative median of 4 (IQR: 1 - 597), followed by nursery farms with 2 (IQR: 0 - 201), sow farms with 1 (IQR: 0 - 132) and wean-to-finish farms with 1 (IQR: 0 - 64) (Supplementary Material, Figure S3) which corresponds to 0.4% (IQR: <0.1% - 55.4%) of the finisher population, 0.5% (IQR: 0% - 51.7%) of the nursery population, 0.3% (IQR: 0% - 44.6%) of the sow population and 0.6% (IQR: 0% - 36.0%) of the wean-to-finish population. Boar studs, farrow-to-finish, gilt, and isolation farms all had a median of 0 infections, however, boar studs and gilts had an upper interquartile range of two and three, respectively. In the NRP simulations, ASF was eliminated within 12 months in 65.4% of the simulations. For the remaining 34.6% of simulations, where ASF was not eliminated, the cumulative median number of infected farms at the end of the simulation (day 450) was 1,383 (IQR: 1,085 - 1,475).

By day 450, the NRP scenario required the depopulation of 32,078 (IQR: 6,480 - 3,883,995) pigs, which consisted of 17,992 (IQR: 2,880 - 2,272,597) finisher pigs, 7,680 (IQR: 0 - 838,307) nursery pigs, 4,350 (IQR: 0 - 386,910) wean-to-finish pigs, and sow pigs with a total of 2,605 (IQR: 0 - 337,235) (Supplementary Material, Figure S4). Boar studs, farrow-to-finishers, gilts, and isolation pigs had medians of 0 pigs depopulated, though boar studs and gilts had upper interquartile ranges of 283 and 2,000, respectively. Similarly, a significant number of diagnostic tests are required by day 450, with a total of 14,240 (IQR: 3,037 - 114,518) which equated to a median of 288 (IQR: 0 - 762) tests for direct contact farms, 1,959 (IQR: 312 - 4,045) tests for indirect contact farms, 3,498 (IQR: 660 - 66,006) tests for farms in infected zones, 3,584 (IQR: 750 - 21,287) tests for farms in buffer zones, and 4,448 (IQR: 942 - 21,143) tests for farms in surveillance zones (Supplementary Material, Figure S5). Over the 450 days, a median of 213 (IQR: 45 - 13,594) permits were required, predominantly for pig movements which accounted for a median of 106 (IQR: 24 - 1,868) permits, followed by feed trucks with a median of 69 (IQR: 13 - 7,211) permits, undefined trucks with a median of 34 (IQR: 5 - 4,245) permits and market trucks with a median of 5 (IQR: 0 - 117) permits (Supplementary Material, Figure S6), resulting in a median of 3,924 (IQR: 483 - 102,600) pre-permit tests. The contribution of each transmission route to infection in the NRP scenario can be found in the Supplementary Material Figure S7.

### 3.4 Controls required to eliminate ASF in three, six, nine, and twelve months

The results of the control strategies for the three, six, nine, and twelve-month time frames suggest that intensifying the implementation of contact tracing, control areas, surveillance zones, and repopulation effectively increases the elimination success (Figure 2). Control strategies eleven, eight, six and five were selected as the best elimination scenarios for the twelve, nine, six and three month time frames. To eliminate ASF from our study population, all four scenarios required the following changes: i) no repopulation of depopulated farms until the end of the epidemic, ii) a three-fold and twelve-fold increase in depopulation and diagnostic testing capacity, iii) an average time to baseline detection of six days, and iv) three negative tests before farms in the control area could apply for permits (Figure 2). For the additional controls, the 12 month scenario required a 30 day movement standstill, 30 days of quarantine for contact farms, traceback of contacts for 60 days, maintenance of control areas and surveillance zones for 60 days, and the expansion of the buffer zone radius to five km (Figure 2). In the nine month scenario, the infected zone radius was also expanded to five km; in the six month scenario, the contact traceback and control area/surveillance zone maintenance were increased to 90 days, and the infected zone, buffer zone and surveillance zone radii were increased to ten km each (Figure 2). Finally, the three month scenario required an increase of the movement standstill to 90 days and an increase of the infected zone, buffer zone and surveillance zone radii to 20 km each (Figure 2). The median number of cumulative infections (including the initial seeded infection) at three, six, nine and twelve months for the corresponding scenarios was 3 (IQR: 1 - 6), 3 (IQR: 1 - 8), 3 (IQR: 1 - 9) and 3 (IQR: 1 - 10) respectively (Figure 3). Infection results by production type and elimination scenario are presented in the Supplementary Material Table S4. In Supplementary Material Figure S8, we present the last day of infection for each of the controlled simulations in the four elimination scenarios. The figure demonstrates that in the majority of simulations, ASF is eliminated within 90 to 150 days regardless of the control strategy.

**Figure 2.**
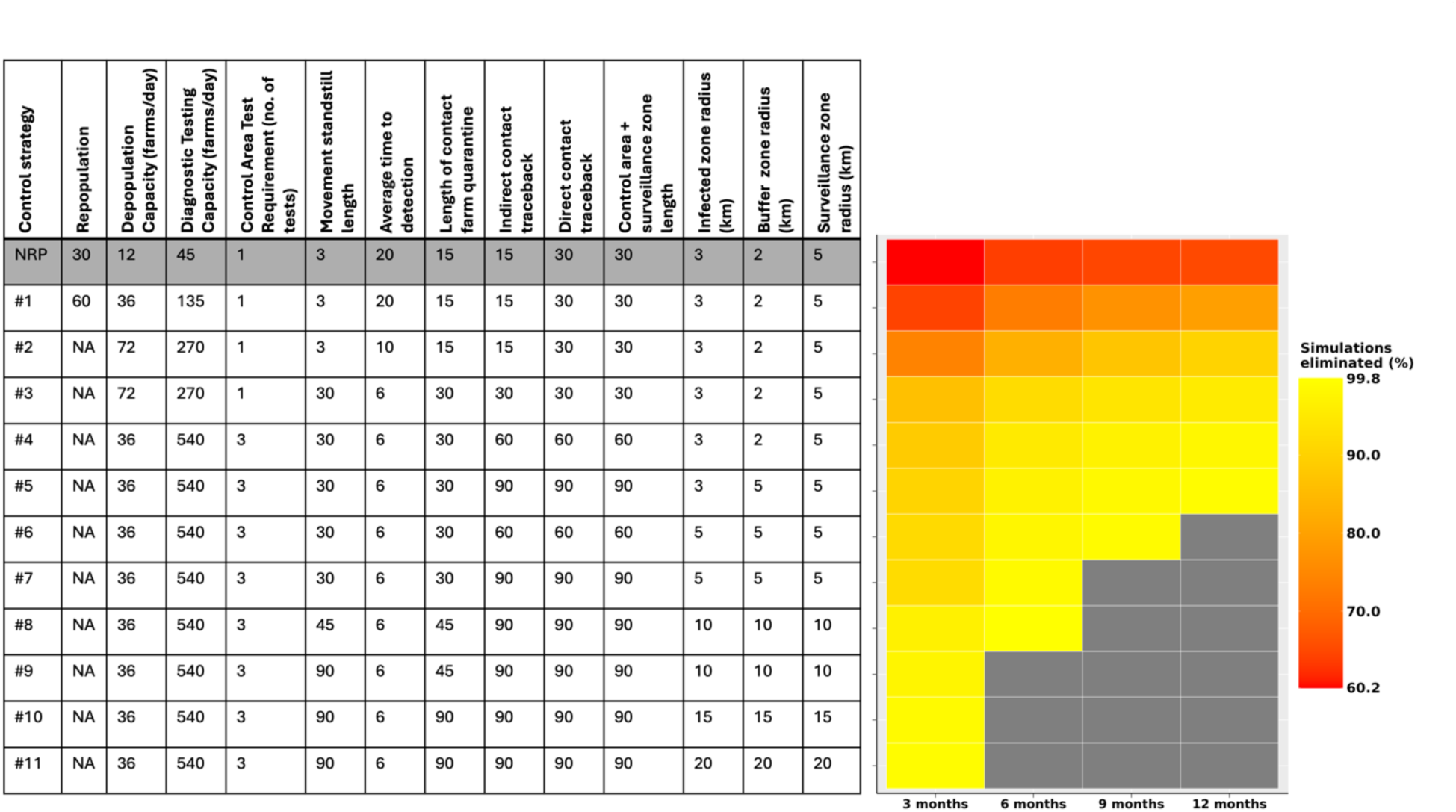
Heat map demonstrating the percentage of simulations where ASF was eliminated within a given time frame. Elimination was determined by a three-month period of no ASF infections following the specified control strategy. The shaded row of the table indicates the control strategy implemented in the NRP scenario. The units are in days unless specified. In the table, “NA” indicates that repopulation was not allowed, while the gray quadrants of the graph indicate that the scenario was not run for that time frame. Elimination percentages are presented in Supplementary Material Table S5.

**Figure 3.**
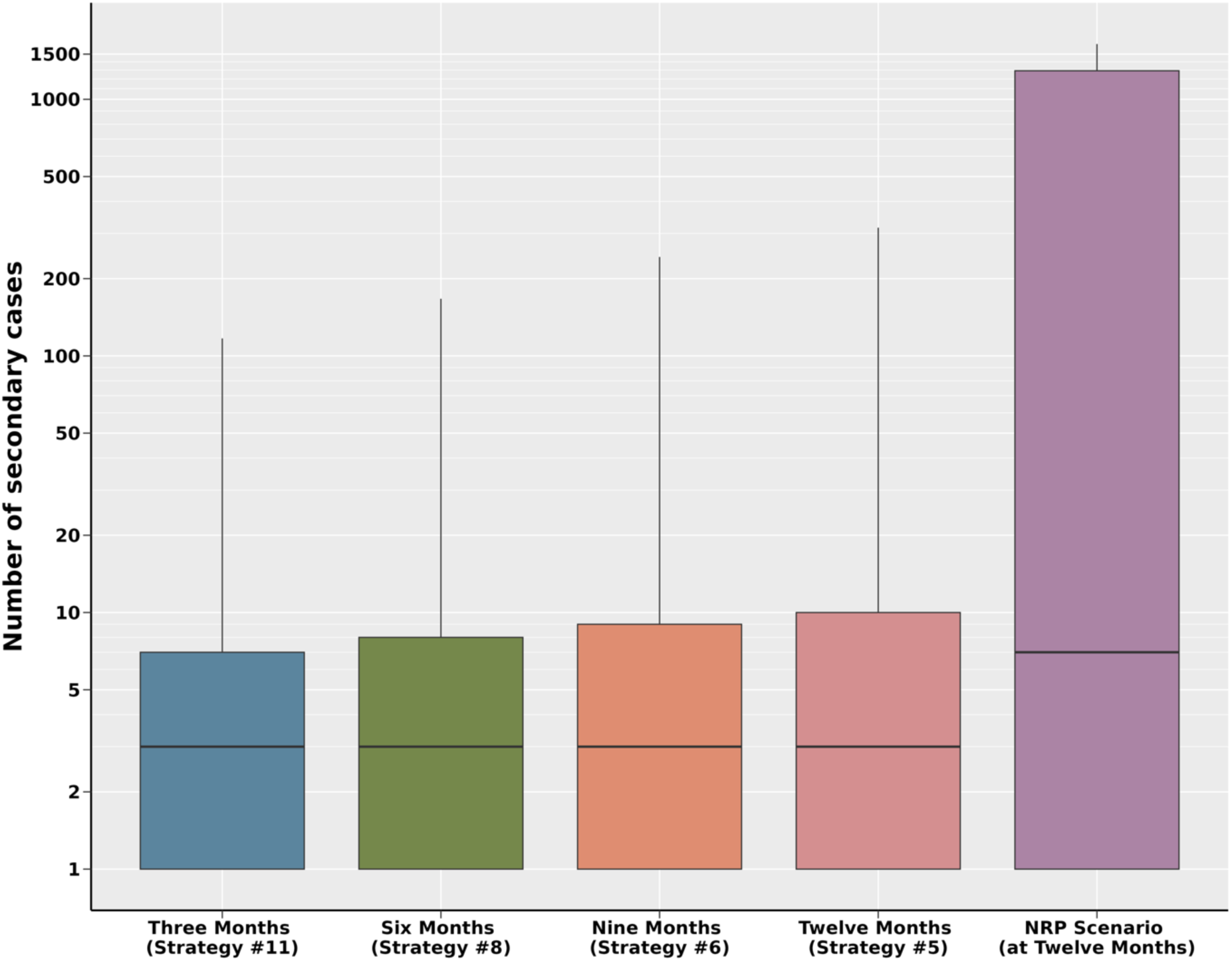
Number of secondary cases in the three, six, nine and twelve month elimination scenarios compared to the NRP scenario at twelve months. The y-axis is shown in log10 format.

### 3.5 Resources required at three, six, nine, and twelve months

For the 12 month elimination scenario, the results show a total of 11,194 (IQR: 4,440 - 45,176) pigs depopulated, which increased to 12,119 (IQR: 4,410 - 42,090) for the nine month scenario, and then decreased to 11,750 (IQR: 4,410 - 35,214) for the six month scenario and 11,520 (IQR: 4,410 - 29,901) for the three month scenario (Figure 4A). For diagnostic tests, the 12 month scenario required a total of 21,791 (IQR: 8,360 - 69,253) tests, which increased as the time frame reduced, with the nine month scenario requiring 29,730 (IQR: 11,281 - 84,166) tests, the six month scenario requiring 132,005 (IQR: 53,327 - 260,469) tests and the three month scenario requiring 341,562 (IQR: 172,074 - 500,547) tests (Figure 4B). Similar trends were demonstrated for the number of permits and pre-permit tests (Figures 4C and 4D). The 12 month scenario required 256 (IQR: 98 - 828) permits and 1,632 (IQR: 396 - 5,607) pre-permit tests, while the nine month scenario required 369 (IQR: 142 - 1,106) permits and 2,772 (IQR: 744 - 8,484) pre-permit tests, the six month scenario required 1,838 (IQR: 736 - 3,772) permits and 11,178 (IQR: 3,288 - 26,328) pre-permit tests, and the three month scenario required 2,989 (IQR: 1,448 - 4,746) permits and 33,711 (IQR: 12,834 - 63,683) pre-permit tests. Detailed results by production type, test reason and movement type for the above scenarios are presented in the Supplementary Material, Tables S6 to S8.

**Figure 4.**
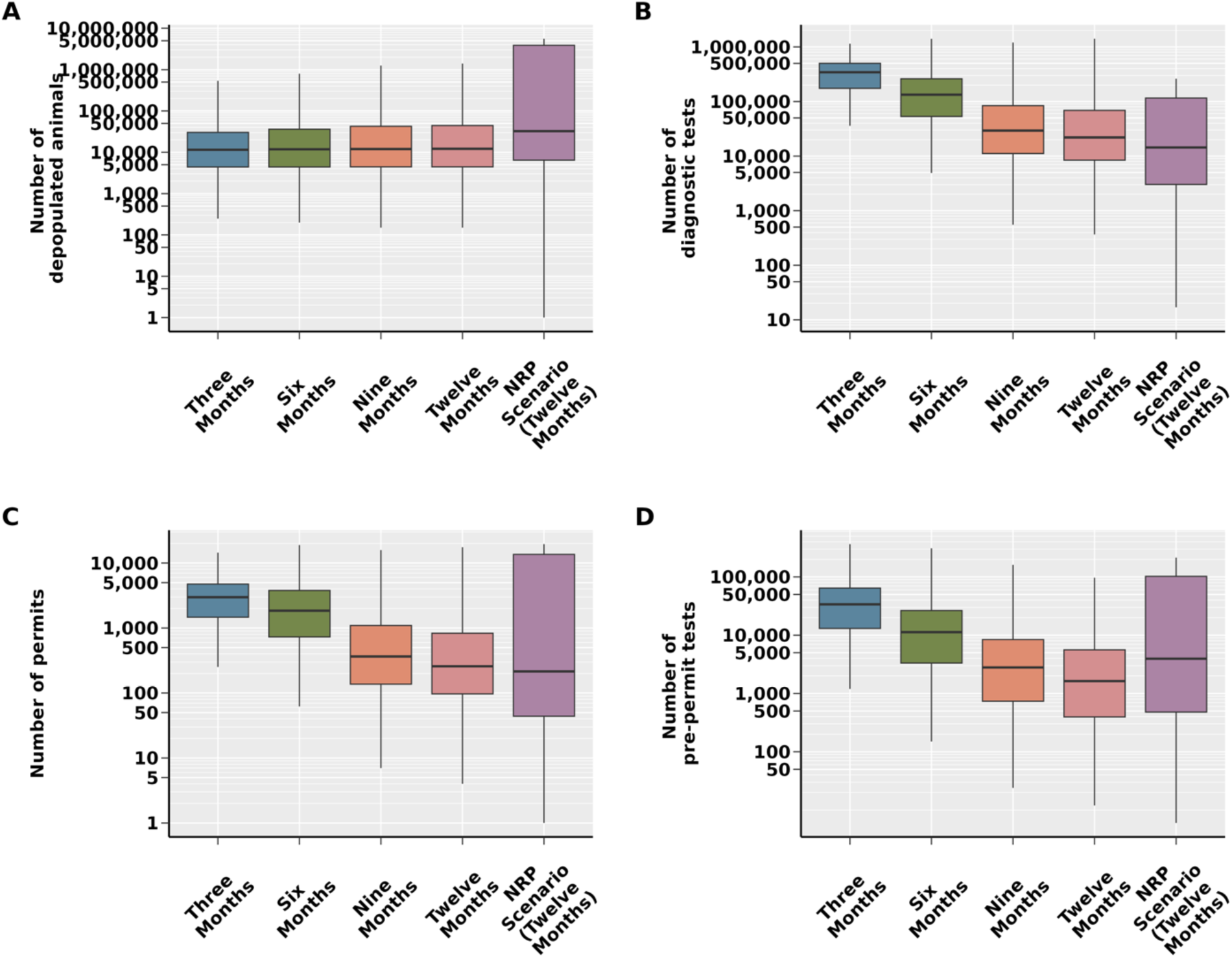
Resource requirements for the three, six, nine and twelve month elimination scenarios in comparison to the NRP scenario. A) The number of depopulated animals; B) the number of diagnostic tests; C) the number of permits; and D) the number of pre-permit tests. The y-axes are shown in log10 format.

### 3.6 Sensitivity analysis

The sensitivity analysis results suggest that the model is most sensitive to changes in parameters that govern the transmission rate of ASF via vehicle movements, the local spread of ASF, surveillance at finisher farms, and the ASF latent period. The PRCC found the linear correlation of the parameters with the number of secondary infections to be statistically significant (Figure 5). More detailed results for the sensitivity analysis are presented in the Supplementary Material Table S9.

**Figure 5.**
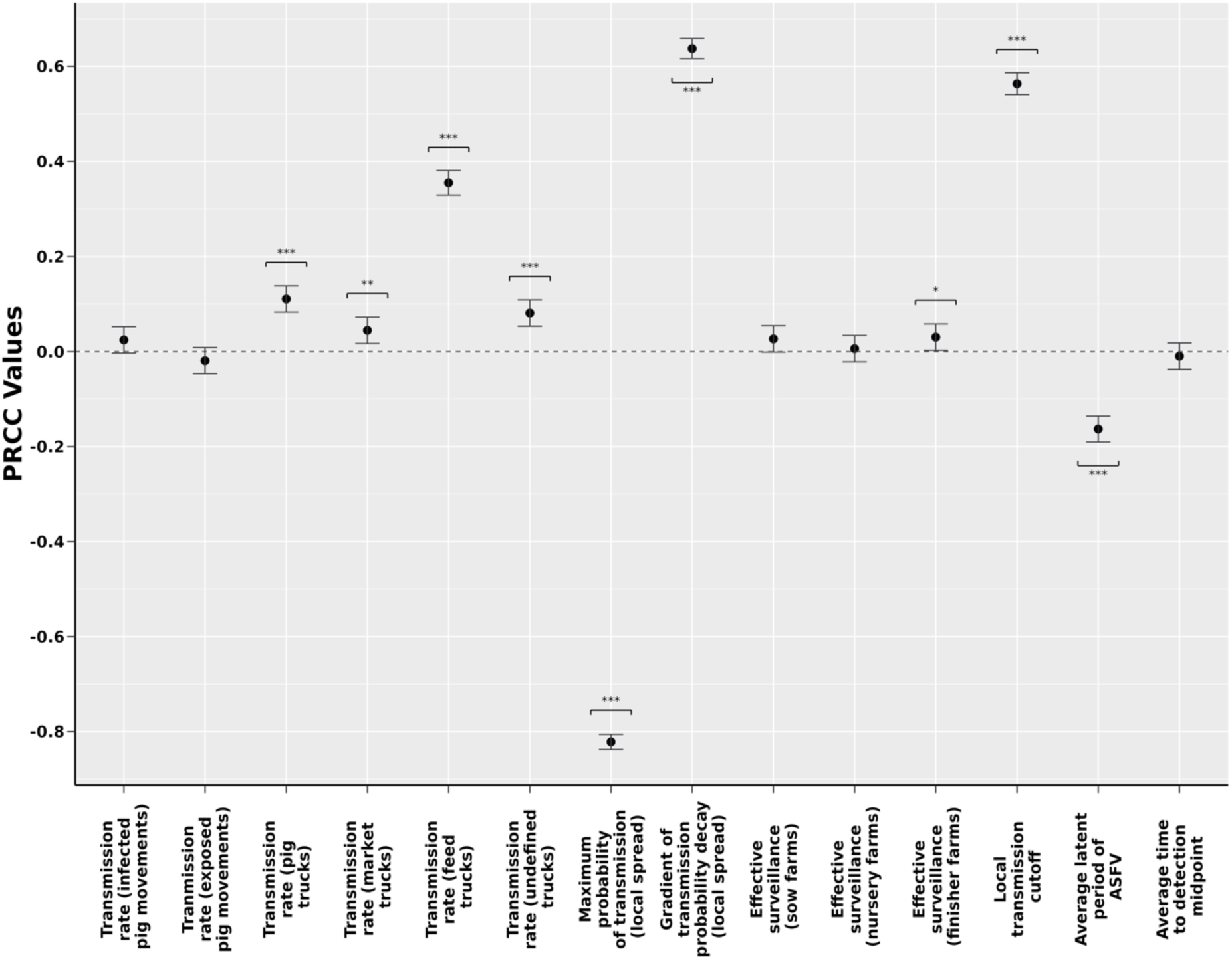
PRCC values indicating the sensitivity of our model to changes in parameters controlling transmission, surveillance, and disease dynamics. The error bars represent 95% confidence intervals. P values: “*” ≤ 0.05, “**” ≤ 0.01, “***” ≤ 0.001.

## 4. Discussion

This study investigated the control strategies necessary to eliminate ASF from the U.S. within three, six, nine, and twelve months by intensifying the control actions listed in the ASF national response plan (USDA, 2023a). Overall, we demonstrated that extending the radii of the infected zones, buffer zones, and surveillance zones up to 20 km each, maintaining control areas for up to 90 days, reducing the time to baseline detection of ASF, and requiring a maximum of 90-day movement restrictions and quarantines effectively reduced the epidemic duration and eliminated between 99 and 99.7% of ASF epidemics within the given time frames. Overall, our results support and guide potential changes to the national response plan to mitigate a lengthy ASF epidemic. Additionally, our model provides estimates of depopulation, diagnostic and permit requirements under different control scenarios, informing resource preparation in anticipation of a future outbreak.

Our elimination scenarios prioritized business continuity, which aimed at minimizing the culling of healthy pigs and restrictions on animal movements, instead focused on increasing diagnostic testing and expanding control areas and surveillance zones (USDA, 2023a). We demonstrated a decrease in median total infections from seven in the NRP scenario to three in all four elimination scenarios. It should be noted that the median number of infections in the NRP is low due to 5,079 (25.6%) simulations in which the initial farm did not cause onward infection; however, total infections ranged from 1 to 1,574 suggesting a wide variation in epidemic trajectory for ASF under the NRP scenario. We observe a narrower range under the elimination scenarios where total infections ranged from 1 to 388.

The most conservative ASF elimination scenario was the 12-month time frame, which required: i) an average time to baseline detection for ASF of six days; ii) a three-fold increase in daily depopulation capacity (36 farms) and 12-fold increase in daily diagnostic testing capacity (540 farms); iii) depopulated farms to remain empty for the course of the epidemic; iv) a 30-day pig and associated vehicle standstill; v) a three-month traceback of direct and indirect contacts, and a subsequent 30-day quarantine; and vi) expansion of the buffer zone to five km alongside a quarantine of control area farms until three-negative tests and maintenance of all three zones for 90 days. Below, for each component of the elimination strategy, we discuss its impacts on disease transmission, the enhancements required for the three, six and nine month eliminations, the feasibility of its implementation, and how it compares to historical eradication efforts.

In all elimination scenarios, it was necessary to reduce the time to baseline detection from the 20 days assumed in the NRP scenario to a much lower six days. This reduced the time that ASF positive farms went undetected, subsequently reducing their transmission potential. Using higher-than-normal mortality rates on farms (Malladi et al., 2022) as the only detection criteria in the absence of controls does not provide enough sensitivity to detect ASF in six days, as it can take four to thirty-six days for ASFv to cause mortality depending on the virulence of the strain (Dixon et al., 2019; Pietschmann et al., 2015; Salguero, 2020; USDA, 2023a). This model does not account for the additional testing being performed under USDA’s Swine Hemorrhagic Fever Integrated Surveillance plan, which includes ASF/CSF testing of veterinary diagnostic laboratory submissions, feral swine surveillance samples, slaughter and aggregation point testing, and testing of higher risk swine populations included those with lower biosecurity, being fed waste and with possible feral swine exposure. Additional detection techniques, such as motion-based video surveillance (Fernández-Carrión et al., 2017), which has been shown to detect ASF up to three days before clinical observations, and lateral flow-based rapid tests (Lu et al., 2020), may also support earlier detection of ASF. A literature review by the European Food Safety Authority (EFSA Panel on Animal Health and Welfare (EFSA AHAW Panel) et al., 2021) described that the median time from infection to detection is 13 days; however, it could be as low as three days, supporting the potential for early detection of ASF in the U.S..

With the intensification of control actions, we also increased the capacity of depopulation and diagnostic testing by three-fold and twelve-fold, from 12 and 45 farms daily in the NRP scenario to 36 and 540 in the 12-month scenario, respectively. This increase in capacity decreases the testing and depopulation delays that have been observed in previous outbreaks of ASF and FMD (Hsu et al., 2023; Nishiura and Omori, 2010), speeding up the identification of undetected infections and reducing the local spread from farms awaiting depopulation. This study used local laboratory capacity to inform baseline for diagnostic testing. The National Animal Health Laboratory Network (NAHLN) currently includes 50 laboratories approved to test for ASF; these laboratories could be activated and provide surge capacity in the event of an outbreak. This work highlights the value and need for this increased capacity to support an efficient response. With this infrastructure in place, the true limitation is the collection and submission of samples, rather than the capacity to analyze them. At the time of writing, such capacities for depopulation and sampling are not attainable (State Animal Health Official, personal communication, 2022; 2023). Currently, the only authorized tissues for live animal ASF diagnostic testing are blood samples, blood swabs and tonsil scrapings (USDA, 2023a), which require varying levels of time and skill to be collected. If oral fluids were to be authorized for an integrated surveillance strategy with blood samples (Galvis et al., 2024; Goonewardene et al., 2021), specifically the more efficient blood swabs (Carlson et al., 2018) the required diagnostic sampling capacity may be significantly reduced. This study calculated surveillance and pre-permit movement testing separately. During the highly pathogenic avian influenza (HPAI) outbreak, surveillance testing has been allowed to contribute to pre-permit testing and vice versa. Taking a similar approach for ASF could also reduce the number of tests required.

Across all elimination time-frame scenarios, a ban on the repopulation of depopulated farms until the end of the epidemic was required. Prohibiting repopulation reduces the number of susceptible farms in the area, lowering the potential for onward transmission, and in a real outbreak, would reduce the possibility of re-infection of a herd from the farm environment. With a median of three total infections across all the elimination scenarios, it could be feasible to consider such strict requirements for repopulation due to the minimal impact on business continuity. Approaches to repopulating farms in past eradication programs have been conservative; sentinel pigs were placed in farms before complete repopulation in Spain, Brazil, and Haiti (Alexander, 1992; Arias and Sánchez-Vizcaíno, 2002; Bech-Nielsen et al., 1995; Danzetta et al., 2020); Italy and Brazil required farms to be empty for six months before repopulation (Danzetta et al., 2020; Lyra, 2006).

In the 12-month elimination scenario, we extended the movement standstill to 30 days, a substantial increase compared to the three-day standstill implemented in the NRP scenario. This was increased to 90 days for the three-month scenario. A lengthy movement standstill is beneficial as it can limit the spread of ASF through live animal movements and associated vehicles (USDA, 2023a; Yoo et al., 2020). However, it greatly impacts business continuity as pigs cannot be sent to slaughter (USDA, 2023a). Producers and swine companies would face issues with overcrowding and feed interruption in as little as two weeks (Weng, 2016), resulting in animal welfare concerns. Financial ramifications would arise from the low cost-benefit of feeding pigs above a certain weight (National Pork Board, 2023). Additionally, the feeding of pigs beyond typical markets weights could result in further downstream complications upon release of the standstill due to the weight limits applied to commercial swine in some processing plants (Corkery and Yaffe-Bellany, 2020), leading to penalties or refusal to slaughter the animal. The financial impact of a long movement standstill could be mitigated by allowing movements to slaughterhouses with high biosecurity to limit the spread of ASF, as discussed in the USDA’s Red Book (USDA, 2023a) and Danzetta et al. (2020).

Additionally, in the 12-month elimination scenario, we increased the traceback of contacts of detected farms (via pig and vehicle movements) to 60 days prior to the date of detection in the herd and required a 30-day quarantine for any contacts found. In the six month scenario, the traceback was extended to 90 days with a 30 day quarantine, and in the three month scenario the quarantine duration was also increased to 90 days. By extending the traceback time we increase the detection of farms which may have been infected for a long period of time without detection, reducing overall onward transmission. This is important as our study population is highly connected (Cardenas et al., 2024), with a median of 91 and 158 movements of pigs and vehicles per day; however, increasing the intensity of contact tracing will require additional personnel and testing resources, and will significantly expand the number of farms under restrictions (Galvis et al., 2024). In previous ASF eradication efforts, explicit diagnostic testing of contact traces was not implemented as a standard control action, with only Belgium subjecting contact farms to serological/diagnostic testing (Danzetta et al., 2020).

Extending the radius of the buffer zone to five km, combined with a quarantine of control area farms for three negative tests, was necessary to eliminate ASF within 12 months. The infected zone radius was also increased to five km for the nine month scenario, while all three zones were increased to ten km and 20 km for the six and three month scenarios, respectively.

The significant impact of enlarging control areas aligns with the sensitivity analysis, which revealed that slight changes in local spread parameterization could result in marked changes in outbreak trajectory, especially for the NRP model (Supplemental Material Figure 7); this reinforces the need to eliminate local pathways for disease spread through rigorous biosecurity practices and elimination of shared personnel and equipment. The increase in coverage of testing and movement restrictions under the expanded control area zones increases the identification of ASF positive farms and reduces the transmission potential within the area, as observed in the Danish ASF model (Halasa et al., 2018, 2015). Alas, the increase in diagnostic samples that would need to be collected with larger control areas and surveillance zones is likely to overwhelm response personnel, making its implementation unfeasible at the time of writing (State Animal Health Officials, personal communication, 2022, 2023)(Supplementary Material, Table S10). This highlights the value of programs such as the certified swine sample collector (CSSC) program that aims to train additional personnel to collect samples in the events of an outbreak. Previous eradication programs in Spain, Italy, and Cuba have implemented area-based restrictions with varying coverage (Arias and Sánchez-Vizcaíno, 2002; Bech-Nielsen et al., 1995; Danzetta et al., 2020; Simeón-Negrín and Frías-Lepoureau, 2008). Spain implemented a three km protection zone and a ten km surveillance zone (Arias and Sánchez-Vizcaíno, 2002; Bech-Nielsen et al., 1995), while Italy placed Rome and its entire metropolitan area into protection and surveillance zones (Danzetta et al., 2020). Cuba did not explicitly place zones on surrounding infected farms but enforced strict quarantines at the province level (Simeón-Negrín and Frías-Lepoureau, 2008).

Overall, our results demonstrate that with an intensification of the national response plan and increases in available sample collection, testing, and depopulation resources, it is likely that an ASF epidemic in the domestic swine population could be eliminated within 12 months. In contrast with smaller countries such as Cuba and Malta, which have eliminated ASF in a short time frame (Danzetta et al., 2020; Simeón-Negrín and Frías-Lepoureau, 2008), countries with substantial swine industries like Spain and Brazil have experienced prolonged ASF epidemics which lasted from several years to decades (Arias and Sánchez-Vizcaíno, 2002; Danzetta et al., 2020; Lyra, 2006; Moura et al., 2010; Mur et al., 2012). Additional challenges that the U.S. may experience include the wide distribution of feral swine (Brown et al., 2024), the presence of soft ticks (Wormington et al., 2019), and environmental conditions that favor ASF persistence (Blome et al., 2024; Carlson et al., 2020; Davies et al., 2017; Sindryakova et al., 2016). While not explicitly incorporated into this model, feral swine are present in swine-producing areas of the U.S. and are a significant concern for the spread of ASF (European Food Safety Authority (EFSA) et al., 2024; Wormington et al., 2019; Yoo et al., 2021). Control actions directly aimed at the feral swine populations will likely be required to eliminate ASF effectively (Dankwa et al., 2022; Muñoz et al., 2022). While soft ticks were a significant component of ASF spread in Portugal, Spain, and Sardinia, there is sparse information about their potential role in a U.S. epidemic. *Ornithodoros spp* that have demonstrated competence to transmit ASF in laboratory studies are present in the U.S., overlapping with the presence of both domestic and feral swine. Additionally, work is needed to define the disease maintenance and transmission risk posed by these potential vectors. Finally, the ability of ASF to persist for long periods of time in the environment can cause infection in pigs without direct contact with an infected animal.

Experimental research has shown that ASF can remain infectious for up to 8.5 days in fecal matter, 15.3 days in urine, 7 days in soil, 30 days in feed, 60 days in water, and 28 days in bedding materials (e.g., bark) depending on the ambient temperature (Blome et al., 2024; Carlson et al., 2020; Davies et al., 2017; Sindryakova et al., 2016). To avoid environmental infection, strict cleaning and disinfection protocols must be considered (Beato et al., 2022; De Lorenzi et al., 2020; USDA, 2023a, 2023b), and can be complemented by the use of sentinel pigs to ensure cleaning and disinfection was effective (Alexander, 1992; Arias and Sánchez-Vizcaíno, 2002; Bech-Nielsen et al., 1995; Danzetta et al., 2020).

## 5. Limitations and future remarks

While our revised ASF transmission model highlights the control strategies required to eliminate ASF within 12 months, there are model assumptions and data limitations that must be acknowledged. Transmission of ASF via vehicle movements is modeled using GPS data that places a vehicle near a cleaning and disinfection (C&D) station; however, proximity does not guarantee that the vehicle was disinfected (Galvis and Machado, 2023). Additionally, in the vehicle network the effectiveness of visits to C&D stations against ASF transmission is assumed to be all-or-nothing, meaning moderately effective C&D protocols that do not meet the required threshold will have no reduction in virus transmission. In the future, we should re-evaluate the all-or-nothing assumption within the network and instead allow C&D to proportionally reduce virus transmission based on the cleaning effectiveness value assigned in the network (Galvis et al., 2024).

Our model does not account for the roles that smallholder independent swine producers nor does it explicitly model ASF spillover from feral swine (Boklund et al., 2020; Costard et al., 2015). Smallholder farms often have limited biosecurity and lack comprehensive knowledge of ASF, increasing their chances of becoming infected and impacting nearby farms, though they often have fewer direct contacts with other premises (Costard et al., 2015; Muñoz-Gómez et al., 2021). Feral swine are known to be present in our study area (Wormington et al., 2019) and contribute significantly to the risk of ASF infection (Boklund et al., 2020). As data becomes available, the model could be revised to incorporate an explicit force of infection from feral swine through a habitat suitability model (Escobar, 2020; Hayes et al., 2024), and include smallholder and independent farms in the study population.

Limitations are also faced when implementing the control actions. For example, our model did not consider the impacts that farm size, the number of personnel, and the time to reach farms may have on depopulation and diagnostic sampling (Galvis et al., 2024). Assuming that a farm of 5,000 heads of swine could be depopulated or sampled as quickly as a farm with 1,000 heads could lead to overestimating control effectiveness and underestimating resource requirements. Research into the time and personnel constraints on diagnostic testing within the study population has been undertaken by (Galvis et al., 2024). Previous publications have detailed the additional limitations associated with the dataset’s representativeness, compliance, resource capacities, and justification to use PEDV historical outbreaks to calibrate model parameters (Sykes et al., 2023; Galivs et al., 2024) instead of using information from the literature. Ultimately, future interaction of this model shall compare model fitting comparing simulation with PEDV data versus a model with parameters drawn from studies in ASFV infected areas.

## 6. Conclusion

This study demonstrates that intensifying the current control strategy outlined in the USDA’s national response plan would likely eliminate ASF from the U.S. within 12 months of its introduction into the domestic swine population. Increasing the initial movement standstill to 30 days, extending contact tracing to 60 days with a 30 day quarantine, increasing the radii of buffer zones to 5 km, maintaining these zones for 60 days, prohibiting repopulation until the end of the epidemic and improving the time to baseline detection for ASF effectively reduced the duration of a U.S. ASF epidemic with limited impacts on business continuity. However, in our study area, the current available capacity for depopulation (four farms per state per day) and diagnostic sampling (10-25 farms per state per day) is insufficient to support such increases in control implementation. Nevertheless, our study provides beneficial guidance to aid preparation for a future ASF introduction and estimates the infrastructure and personnel required to promptly bring an epidemic under control.

## Footnotes

1 The elimination scenarios are only a recommendation of changes to the USDA’s national response plan, and in no way reflect the opinions or intent of the USDA.

