## Supplementary material for "Identifying control strategies to eliminate African swine fever from the United States swine industry in under 12 months": SS

The following equations represent the dynamics of our *PigSpread* ASF model:

**Transmission routes**

$$Movement\ of\ exposed\ pigs = \lambda_{nit} = \beta_n * N_{nit} \quad (1)$$
$$Movement\ of\ infected\ pigs = \lambda_{sit} = \beta_s * N_{sit} \quad (2)$$
$$Movement\ of\ detected\ pigs = \lambda_{ait} = \beta_a * N_{ait} \quad (3)$$
$$Movement\ of\ pig\ vehicles = \lambda_{pit} = \beta_p * M_{ijt} * Z_{it} \quad (4)$$
$$Movement\ of\ market\ vehicles = \lambda_{mit} = \beta_m * M_{ijt} * Z_{it} \quad (5)$$
$$Movement\ of\ feed\ vehicles = \lambda_{fit} = \beta_f * M_{ijt} * Z_{it} \quad (6)$$
$$Movement\ of\ undefined\ vehicles = \lambda_{bit} = \beta_b * M_{ijt} * Z_{it} \quad (7)$$
$$Local\ spread = \lambda_{lit} = 1 - \prod_1^i \left(1 - \phi e^{-\alpha d_{ij}}\right) \quad (8)$$

**Probability of transmission**

$$Y_{it} = 1 - e^{(-\lambda_{nit} - \lambda_{pit} - \lambda_{mit} - \lambda_{fit} - \lambda_{bit})} + \lambda_{lit} \quad (9)$$
$$V_{it} = 1 - e^{\lambda_{sit}} \quad (10)$$
$$W_{it} = 1 - e^{\lambda_{ait}} \quad (11)$$

**Conditions for transition**

$$Y_{it} > X \sim U(0,1) \quad (12)$$
$$V_{it} > X \sim U(0,1) \quad (13)$$
$$W_{it} > X \sim U(0,1) \quad (14)$$

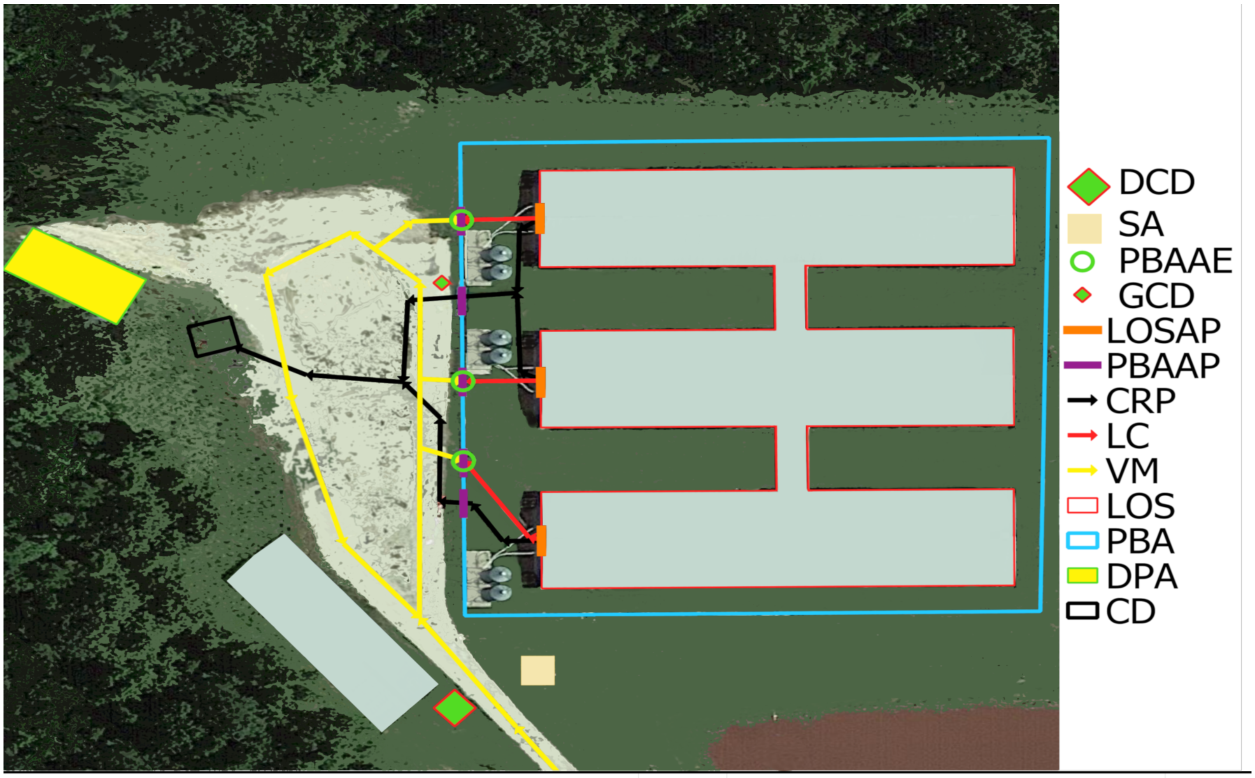


| Legend | Symbol | Legend | Symbol |
|---|---|---|---|
| LOSAE (LOS Animal Emergency) | | PBAAP (PBA Access Point) | |
| SE (Site Entry) | | LOSAP (LOS Access Point) | |
| PBAAE (PBA Access Entry, animals only) | | CRP (Carcasses Removal Pathways) | |
| DCD (Designated Cleaning and Disinfection vehicle station) | | VM (Vehicle Movements) | |
| GCD (Generalized Cleaning and Disinfection temporary station) | | LC (Loading Chute) | |
| PCD (Proposed Cleaning and Disinfection-temporary station) | | PBA (Perimeter Buffer Area) | |
| ADU (Dumpster for dead animals) | | LOS (Line of Separation) | |
| TD (Trash Dumpster) | | DPA (Designated Parking Area) | |
| SA (Supply drop-off Area) | | CD (Carcasses Disposal location/ADU box) | |

**Figure S1. Fabricated farm premises map containing farm features, including the Perimeter Buffer Area (PBA).** This represents an example of the maps developed for farms included in the Secure Pork Supply plans.

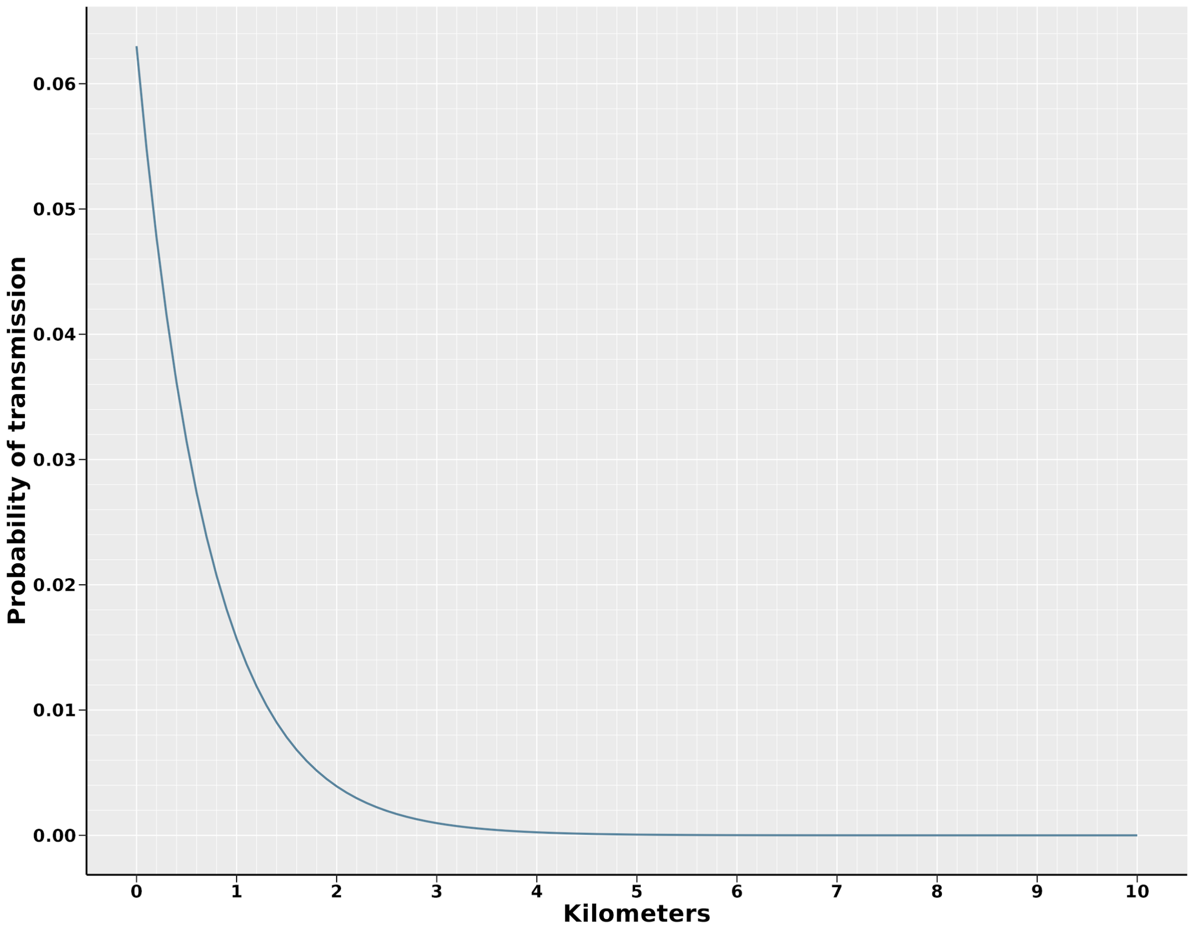


**Figure S2. Probability of local transmission between an infected/detected farm and a susceptible farm over decreasing distances (0 km to 10 km) using the median values for $\alpha$ and $\phi$ produced by the ABC calibration.**

**Table S1. Prior distributions used in the Approximate Bayesian Computation Rejection algorithm to calibrate the model parameters**

| *Parameter* | *Notation* | *Prior Distribution* |
|---|---|---|
| Transmission rate for infected pig movements | $\beta_s$ | Uniform(1.25, 1.75) |
| Transmission rate for exposed pig movements | $\beta_n$ | Uniform(1.25, 1.75) |
| Transmission rate for movements of pig trucks | $\beta_p$ | Uniform(0.01, 0.1) |
| Transmission rate for movements of market trucks | $\beta_m$ | Uniform(0.01, 0.1) |
| Transmission rate for feed delivery trucks | $\beta_f$ | Uniform(0.001, 0.05) |
| Transmission rate for undefined trucks | $\beta_b$ | Uniform(0.01, 0.1) |
| Maximum probability of transmission | $\phi$ | Uniform(0.01, 0.1) |
| Gradient of transmission probability decline over distance | $\alpha$ | Uniform(1.2, 1.6) |
| Effective surveillance in sow farms | $L_{sow}$ | Uniform(0.5, 0.99) |
| Effective surveillance in nursery farms | $L_{nursery}$ | Uniform(0.25, 0.75) |
| Effective surveillance in finisher farms | $L_{finisher}$ | Uniform(0.25, 0.75) |

**Table S2. Testing frequency for contact farms and farms within the infected, buffer and surveillance zones.**

| *Testing reason* | *Frequency of testing* |
|---|---|
| Direct/indirect contact | Every six days until the end of quarantine |
| Infected zone | Two tests at an interval of three days; followed by tests every six days until the zone is lifted |
| Buffer zone | Every six days until the zone is lifted |
| Surveillance zone | Every 15 days until the zone is lifted |

With limited test capacity, testing direct and indirect contacts was prioritized followed by farms in the infected zone, farms in the buffer zone and farms in the surveillance zone

**Table S3. Probability density inputs for the LHS-PRCC sensitivity analysis**

| ***Parameter*** | ***Probability density*** |
|---|---|
| Transmission rate for infected pig movements | Uniform(1, 2) |
| Transmission rate for exposed pig movements | Uniform(1, 2) |
| Transmission rate for movements of pig trucks | Uniform(0.001, 0.1) |
| Transmission rate for movements of market trucks | Uniform(0.001, 0.1) |
| Transmission rate for feed delivery trucks | Uniform(0.001, 0.05) |
| Transmission rate for undefined trucks | Uniform(0.001, 0.1) |
| Maximum probability of transmission | Uniform(0, 1) |
| Gradient of transmission probability decline over distance | Uniform(0.001, 0.1) |
| Effective surveillance in sow farms | Uniform(0, 1) |
| Effective surveillance in nursery farms | Uniform(0, 1) |
| Effective surveillance in finisher farms | Uniform(0, 1) |
| Local transmission spread cut off | Uniform(1, 25) |
| ASF latent period | Uniform(3, 7) |
| Time to detection midpoint | Uniform(5, 20) |

Limitations of LHS-PRCC

While LHS-PRCC is considered an efficient technique for sensitivity analysis; its efficiency and reliability are reduced considerably when dealing with covariances between input parameters or non-monotonic relationships between the input and output parameters (Drake and Rohani, 2016; Marino et al., 2008). Additionally, applying LHS-PRCC to stochastic models, it can be challenging to separate the uncertainty caused by varying the input parameters (epistemic uncertainty) and the uncertainty due to the stochastic nature of the model (aleatory uncertainty) (Marino et al., 2008). To counteract the latter, we seeded the infection on the same farm on the same date for each sensitivity simulation.

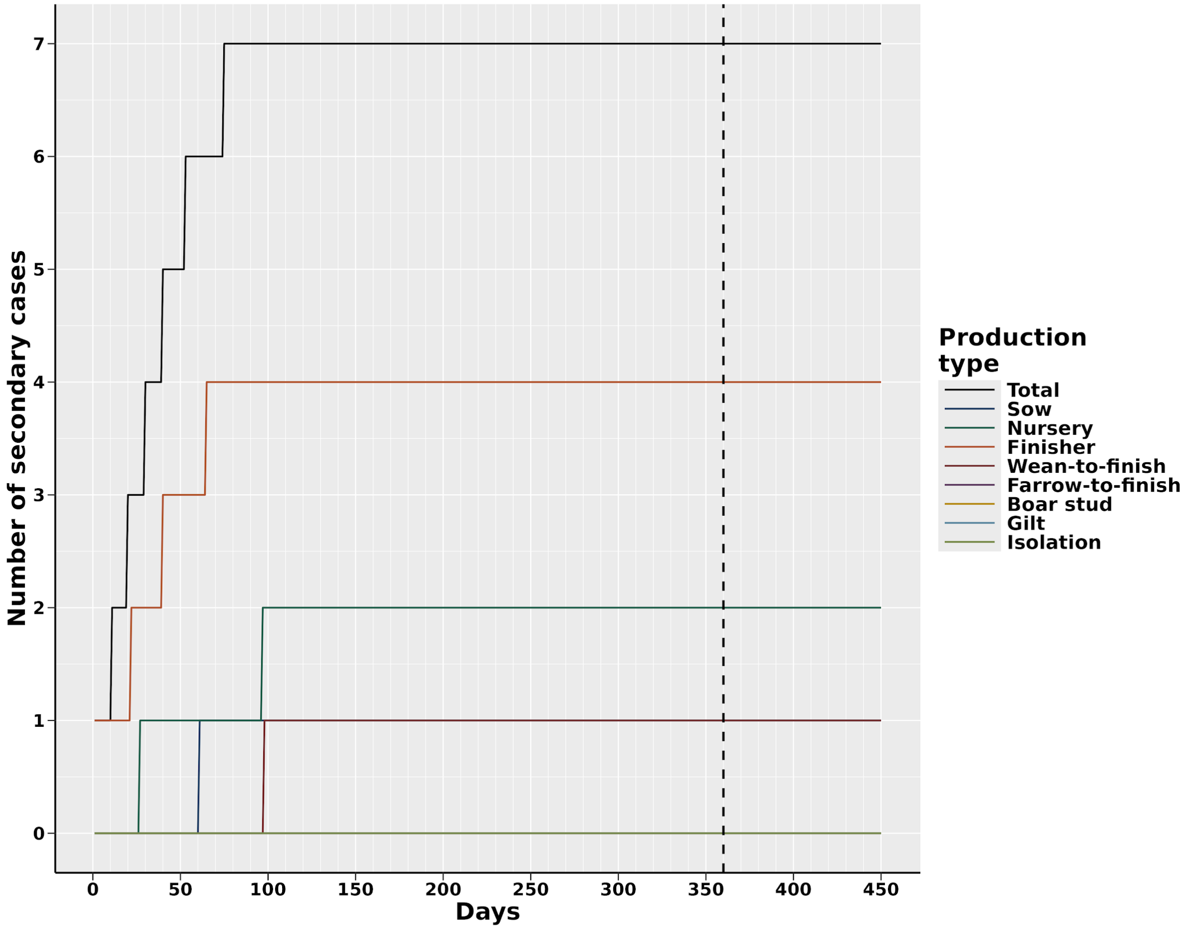


**Figure S3. Median cumulative infections over 450 days of the epidemic, total and by production type, under the NRP scenario.** The dotted line represents the end of the 12-month (360 days) time frame.

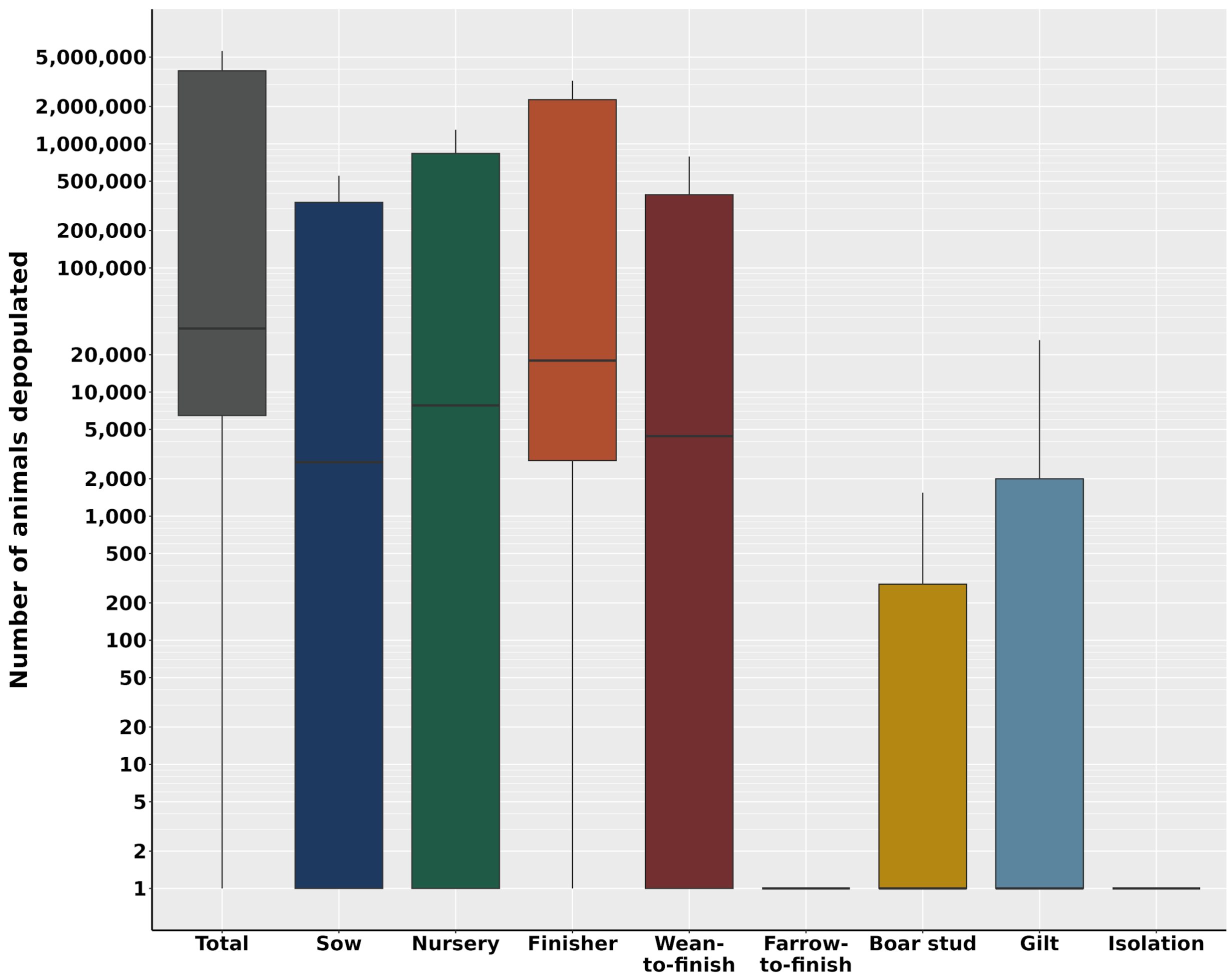


**Figure S4. Median cumulative pigs depopulated by 450 days of the epidemic, total and by production type, under the NRP scenario.**

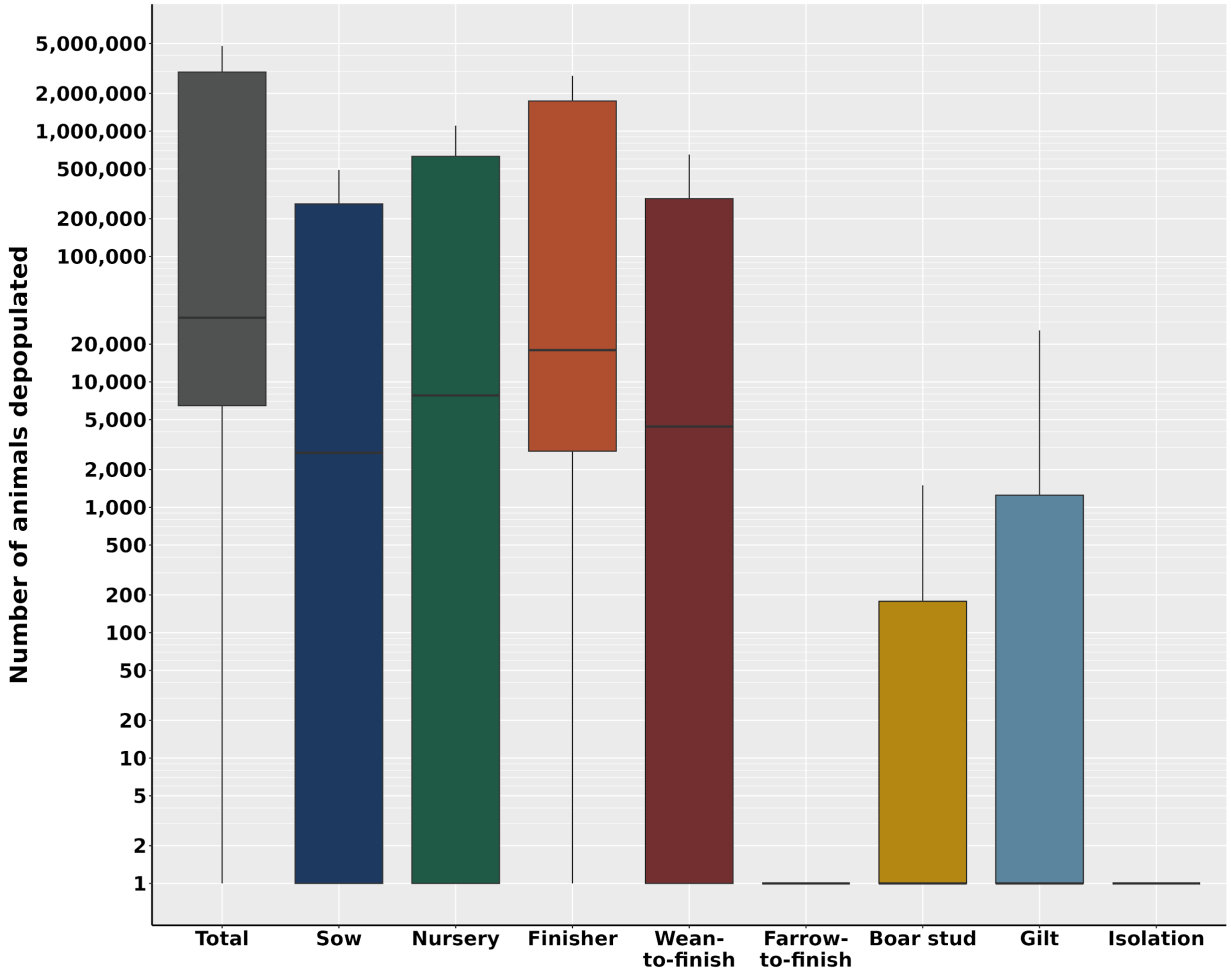


**Figure S5. Median cumulative pigs depopulated by 360 days of the epidemic, total and by production type, under the NRP scenario.**

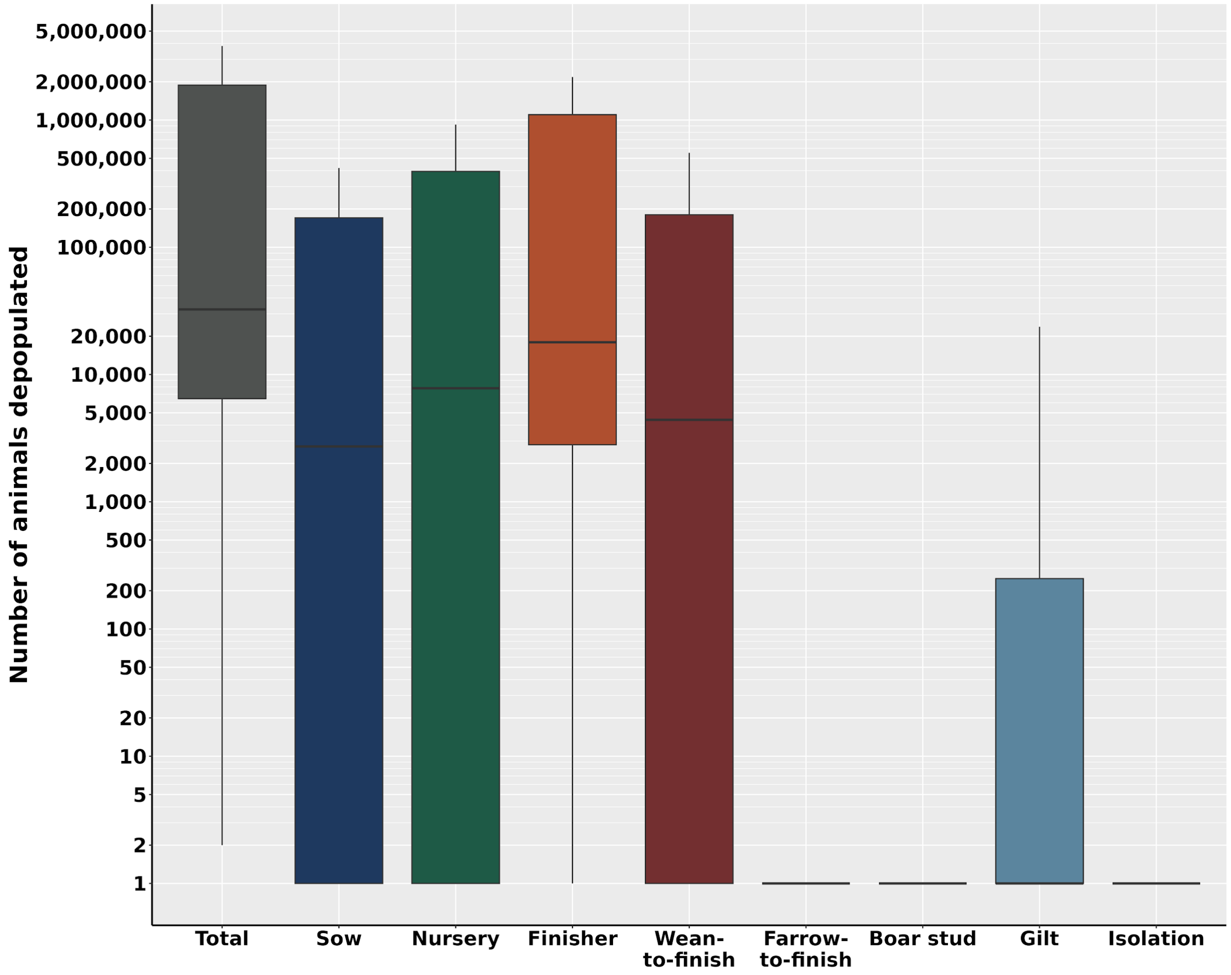


**Figure S6. Median cumulative pigs depopulated by 270 days of the epidemic, total and by production type, under the NRP scenario.**

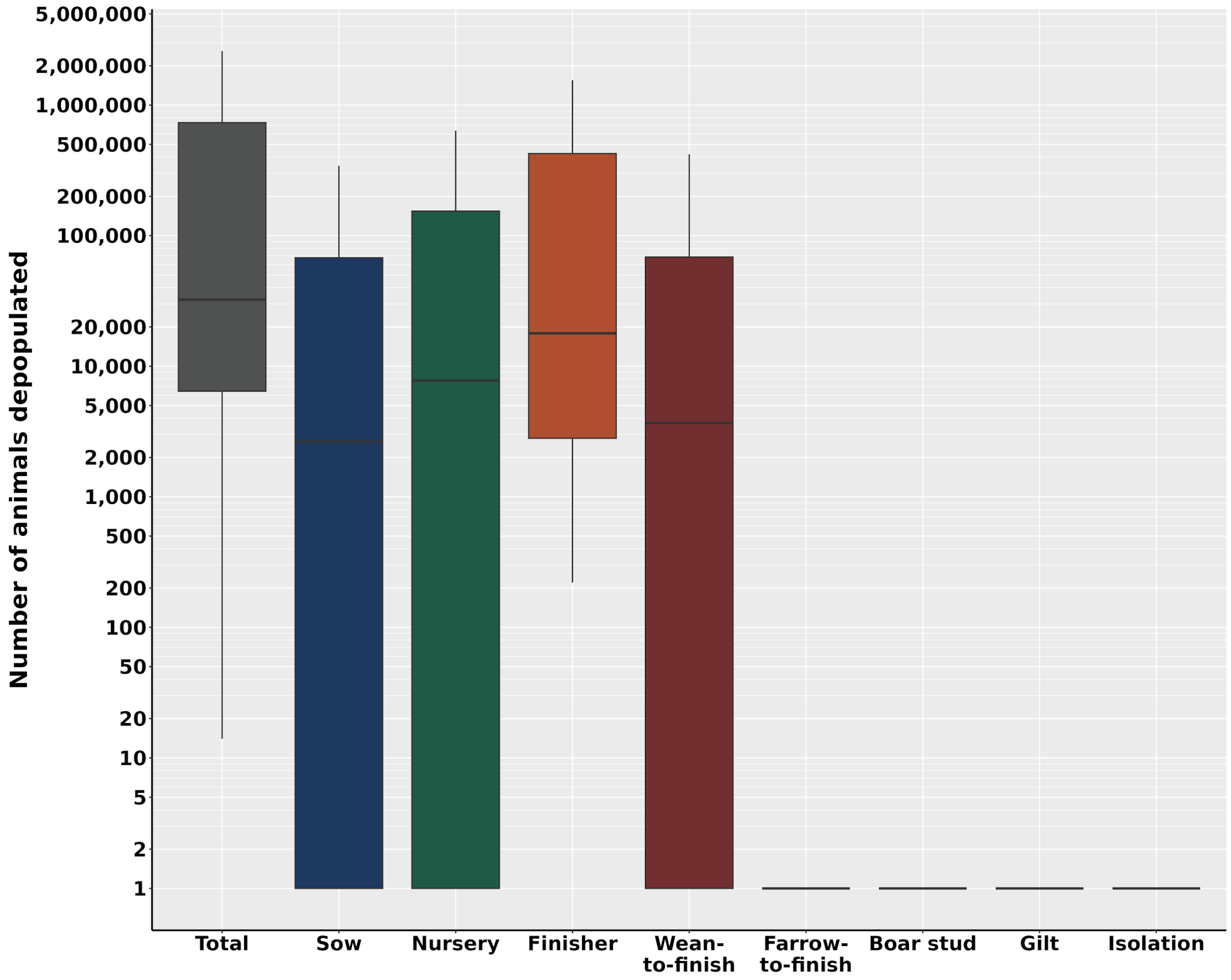


**Figure S7. Median cumulative pigs depopulated by 180 days of the epidemic, total and by production type, under the NRP scenario.**

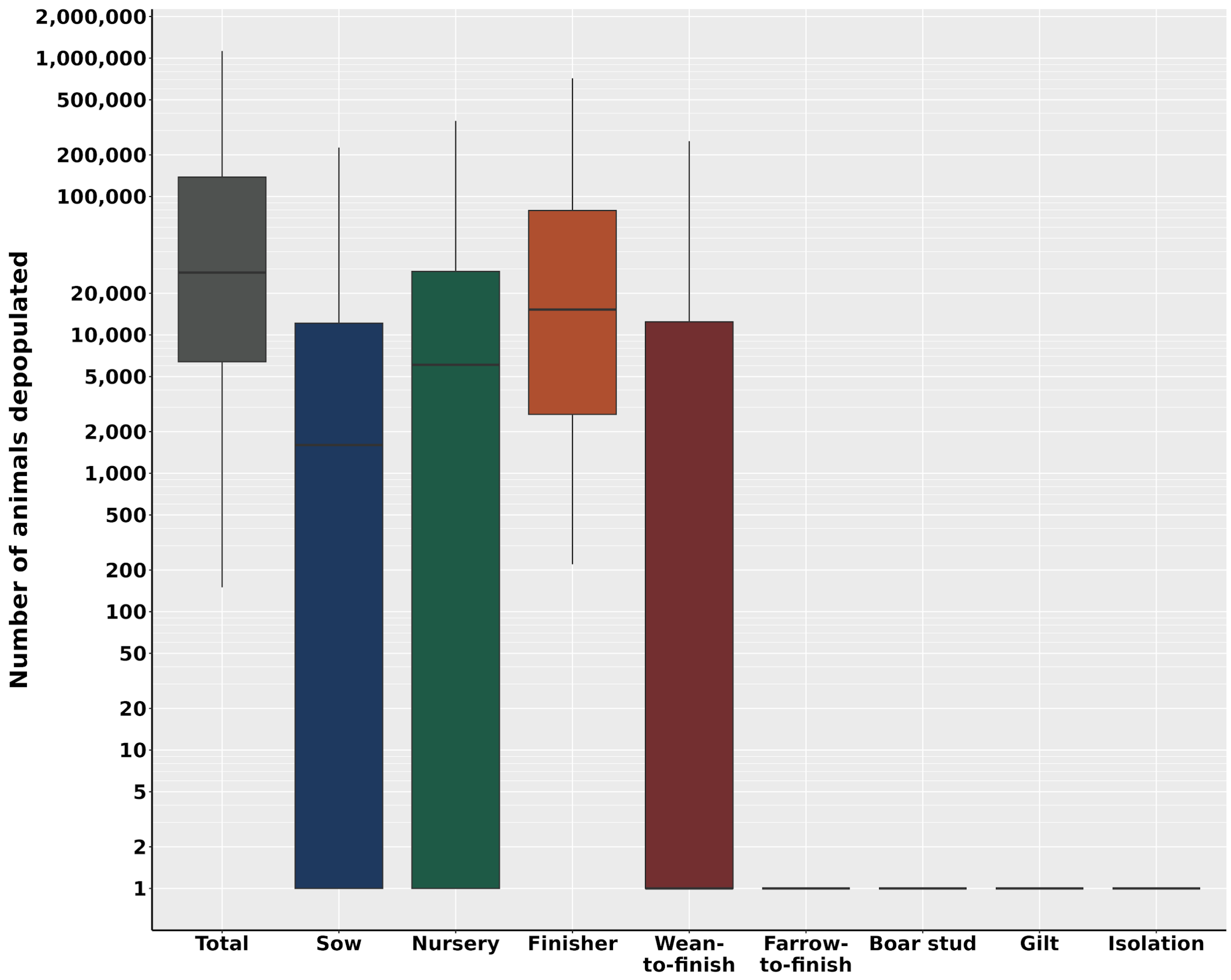


**Figure S8. Median cumulative pigs depopulated by 90 days of the epidemic, total and by production type, under the NRP scenario.**

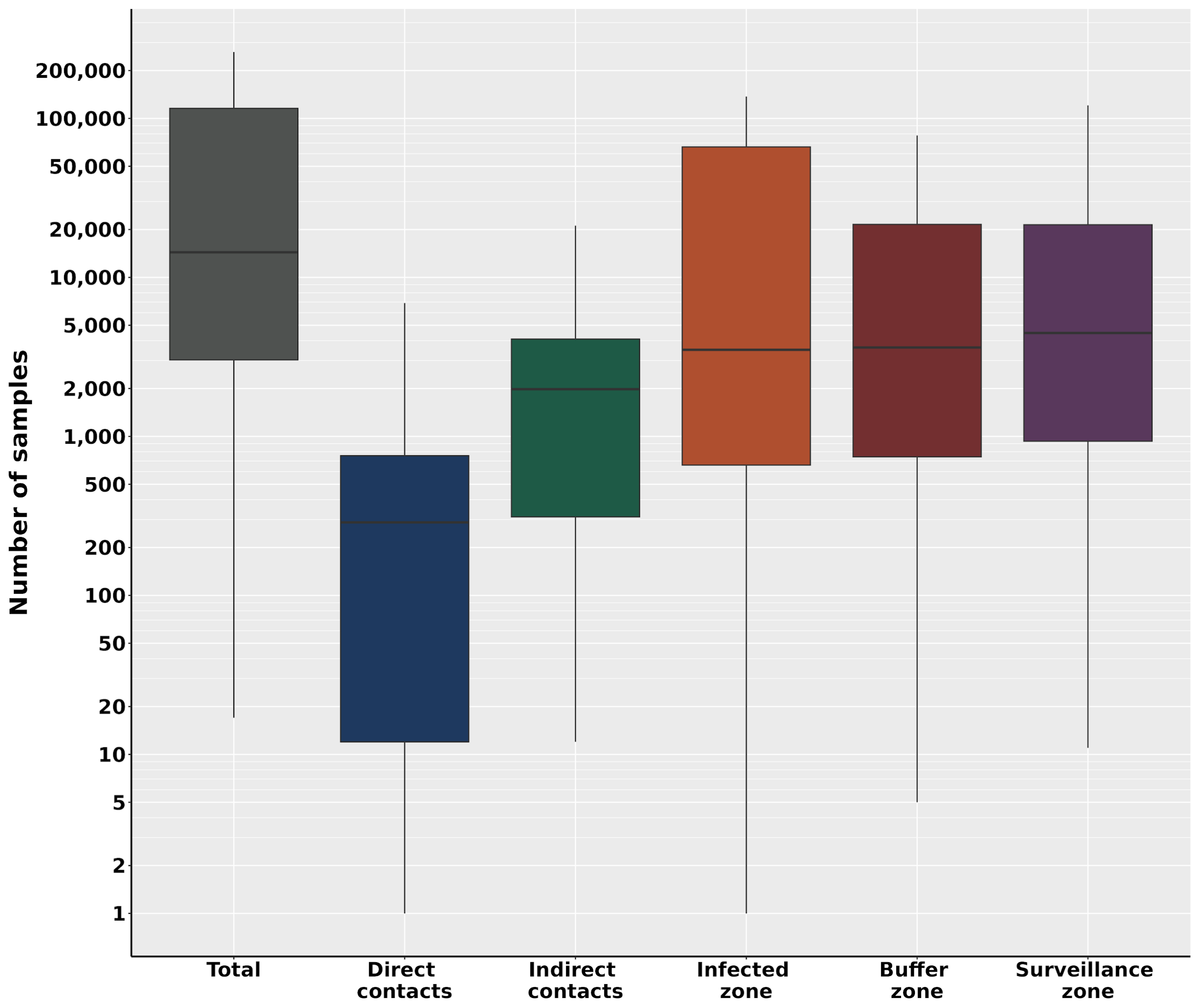


**Figure S9. Median cumulative diagnostic tests required by 450 days of the epidemic, total and by test reason, under the NRP scenario.**

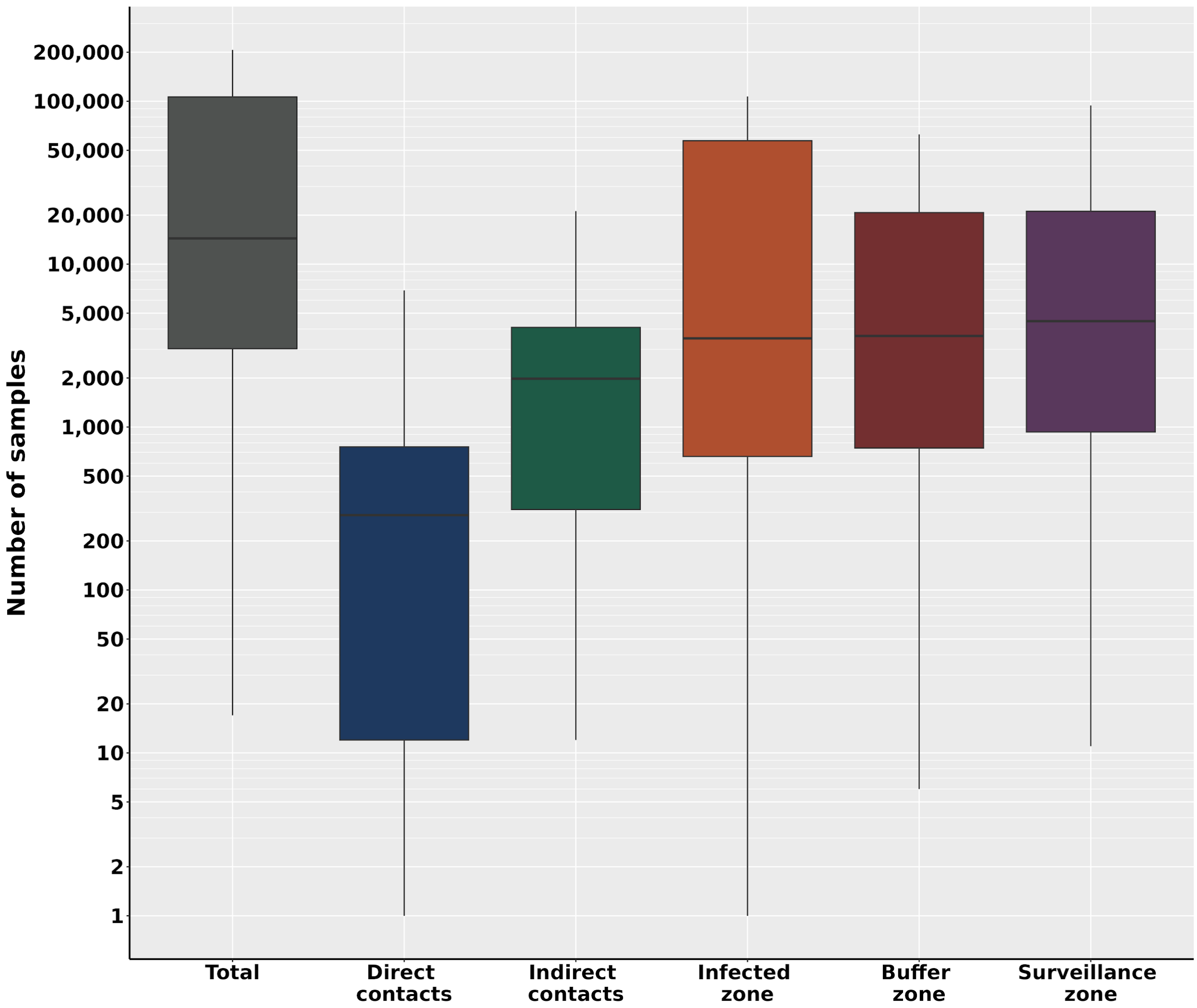


**Figure S10. Median cumulative diagnostic tests required by 360 days of the epidemic, total and by test reason, under the NRP scenario.**

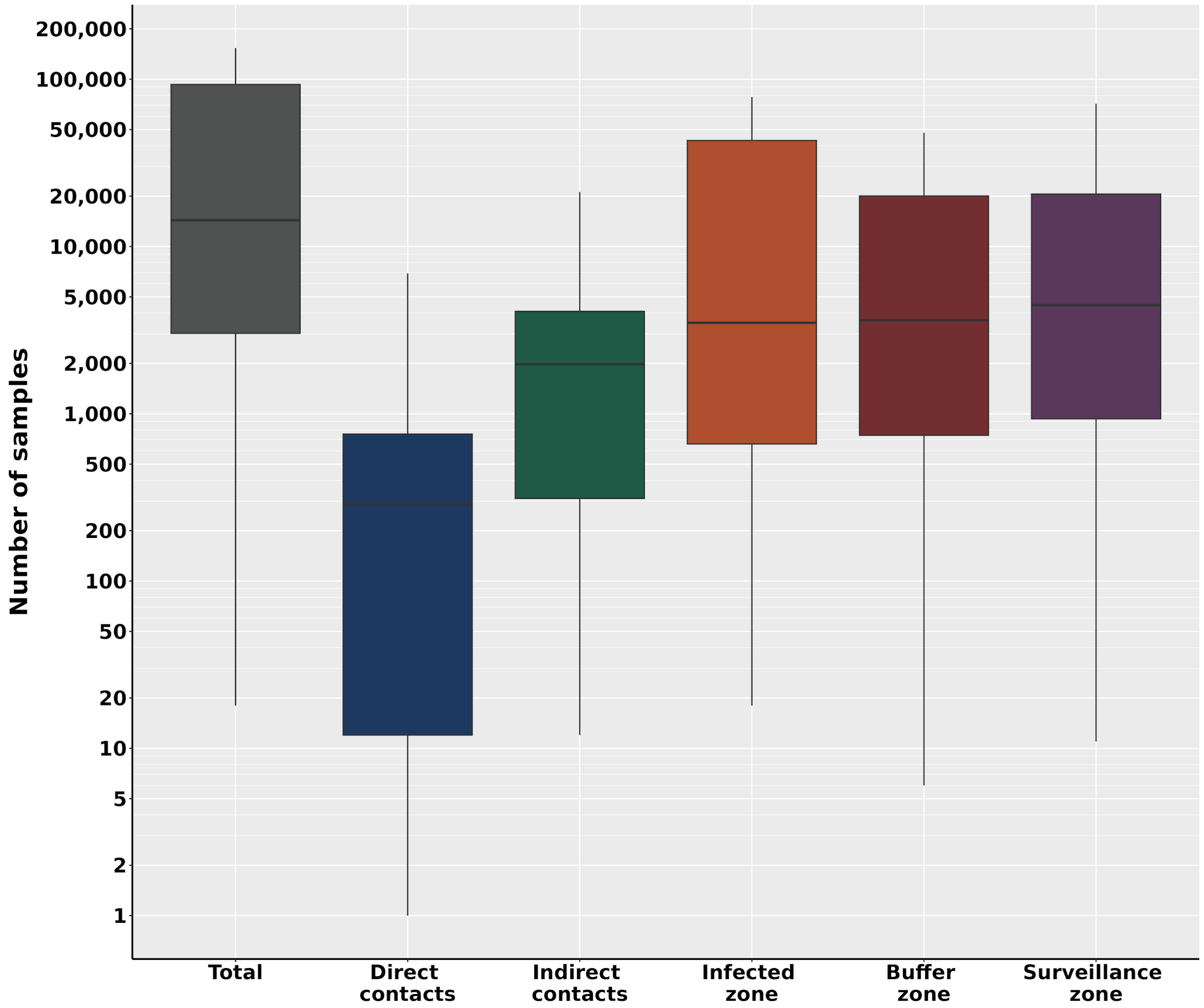


**Figure S11. Median cumulative diagnostic tests required by 270 days of the epidemic, total and by test reason, under the NRP scenario.**

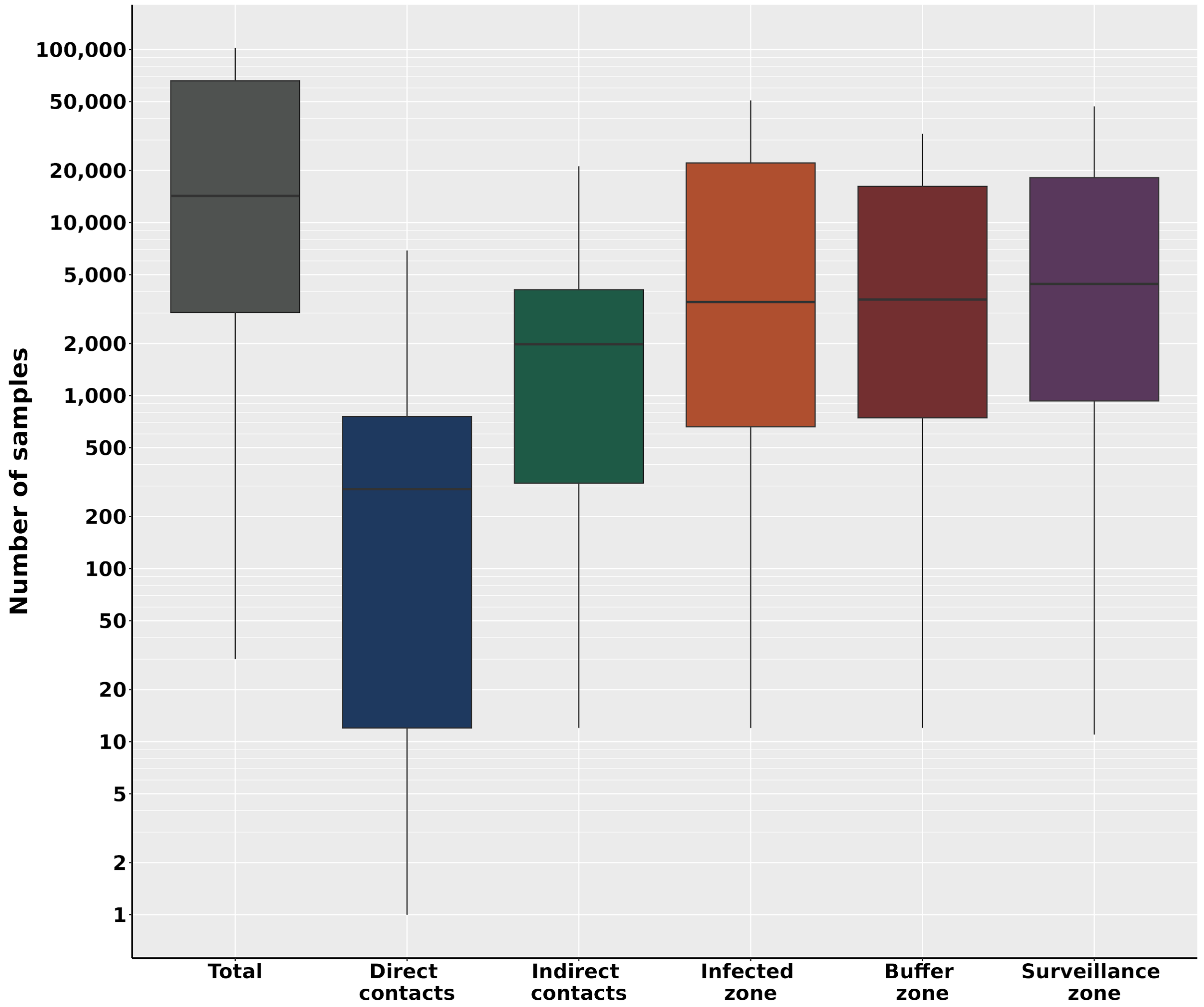


**Figure S12. Median cumulative diagnostic tests required by 180 days of the epidemic, total and by test reason, under the NRP scenario.**

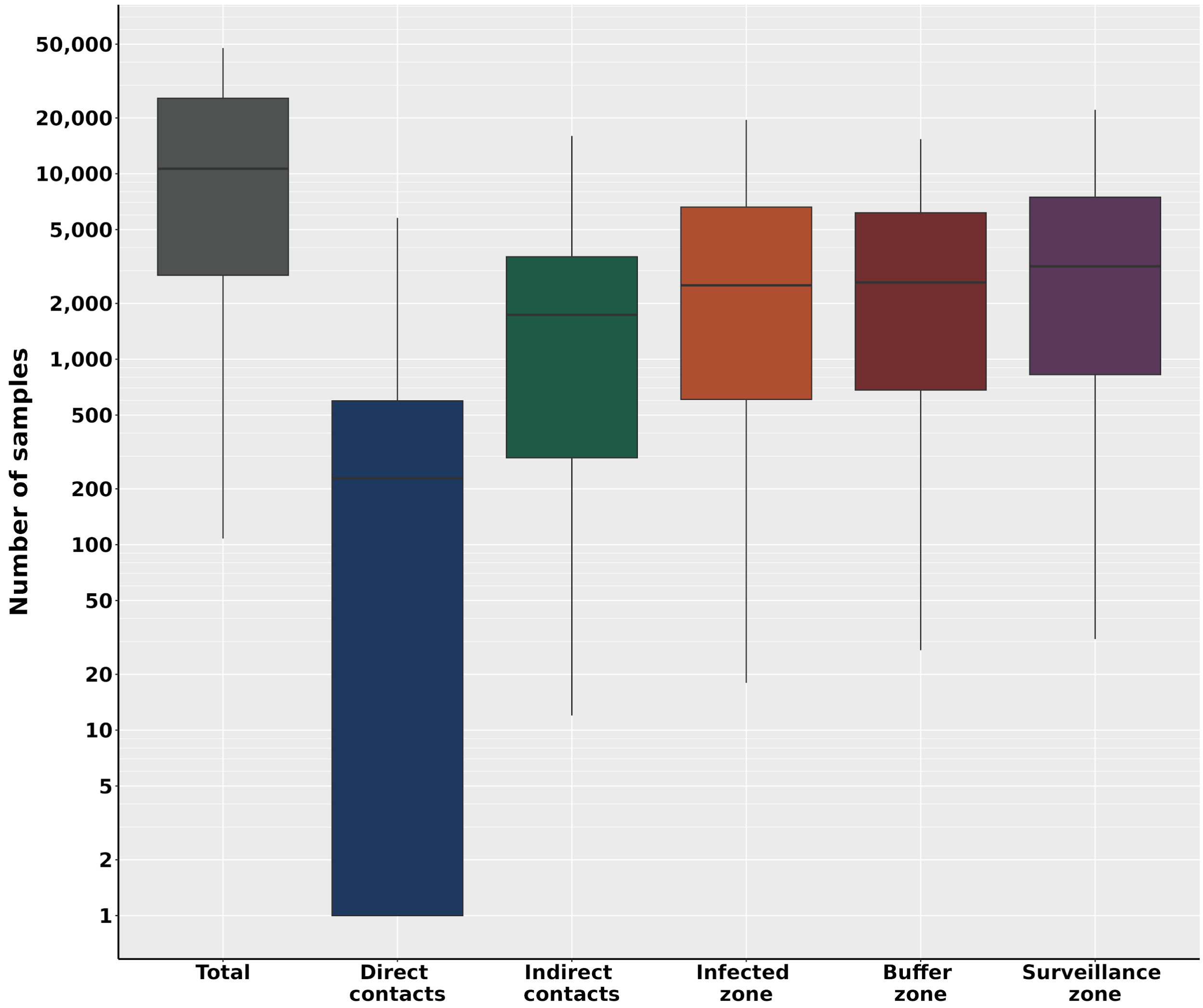


**Figure S13. Median cumulative diagnostic tests required by 90 days of the epidemic, total and by test reason, under the NRP scenario.**

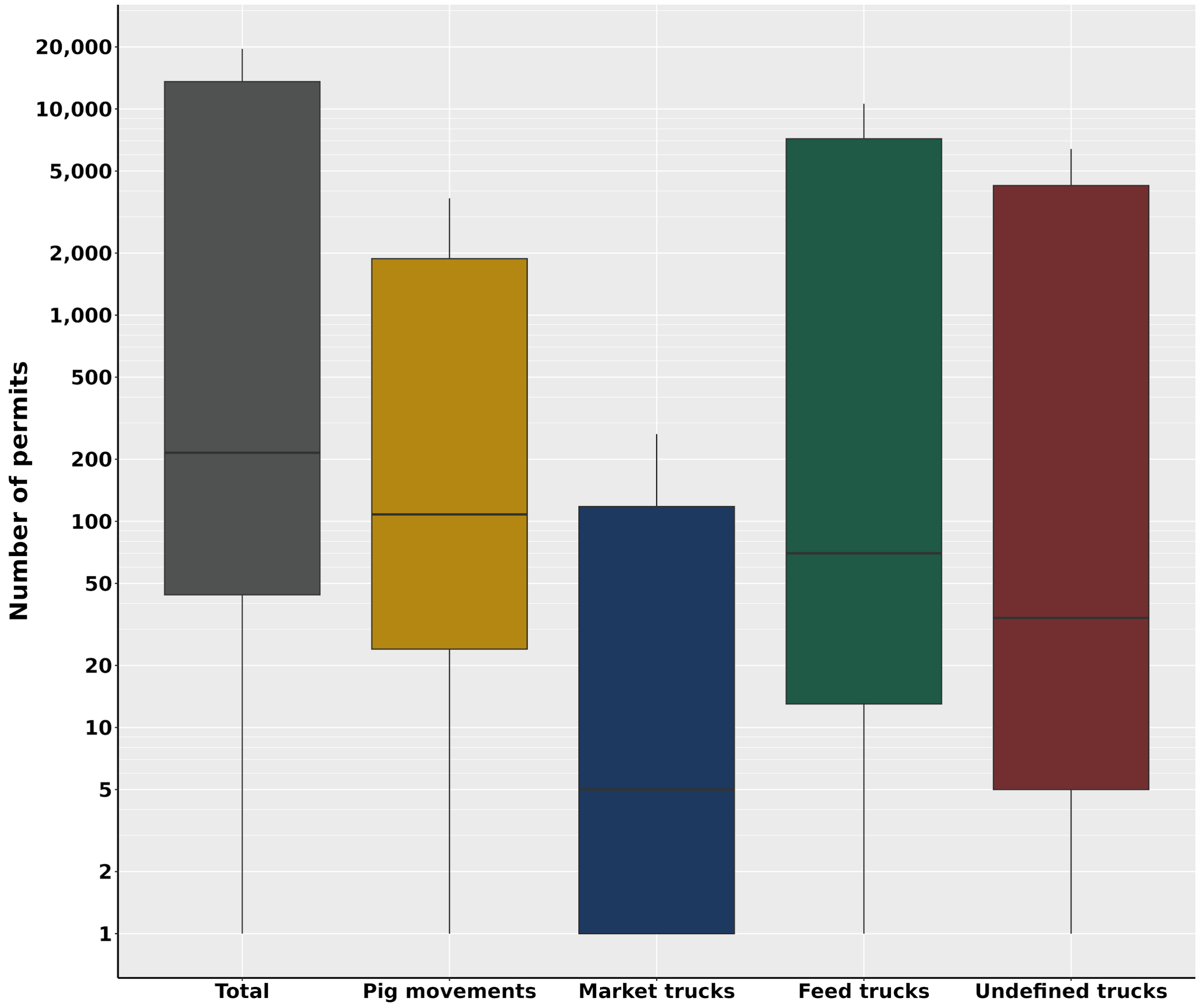


**Figure S14. Median cumulative permits required by 450 days of the epidemic, total and by movement type, under the NRP scenario.**

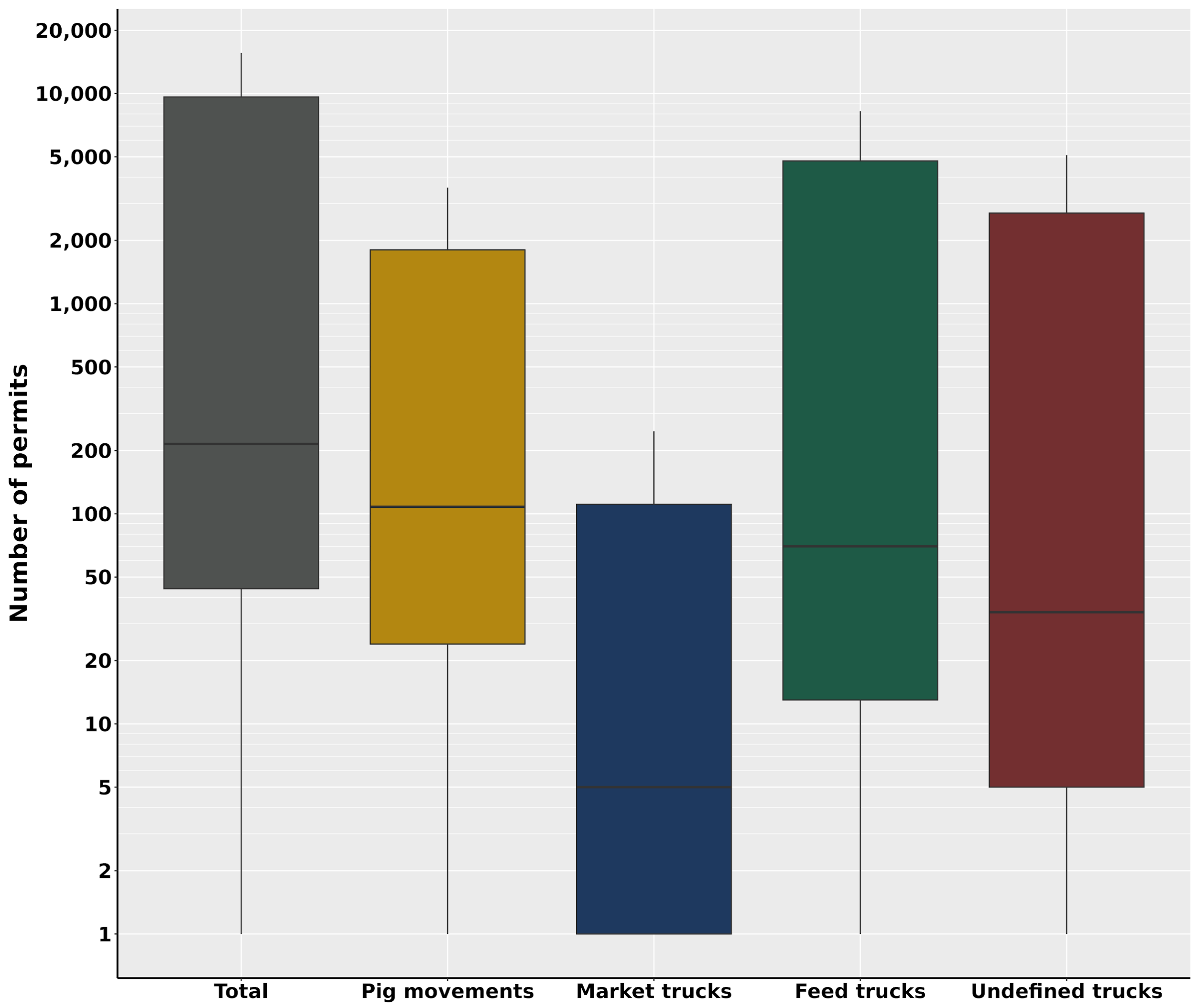


**Figure S15. Median cumulative permits required by 360 days of the epidemic, total and by movement type, under the NRP scenario.**

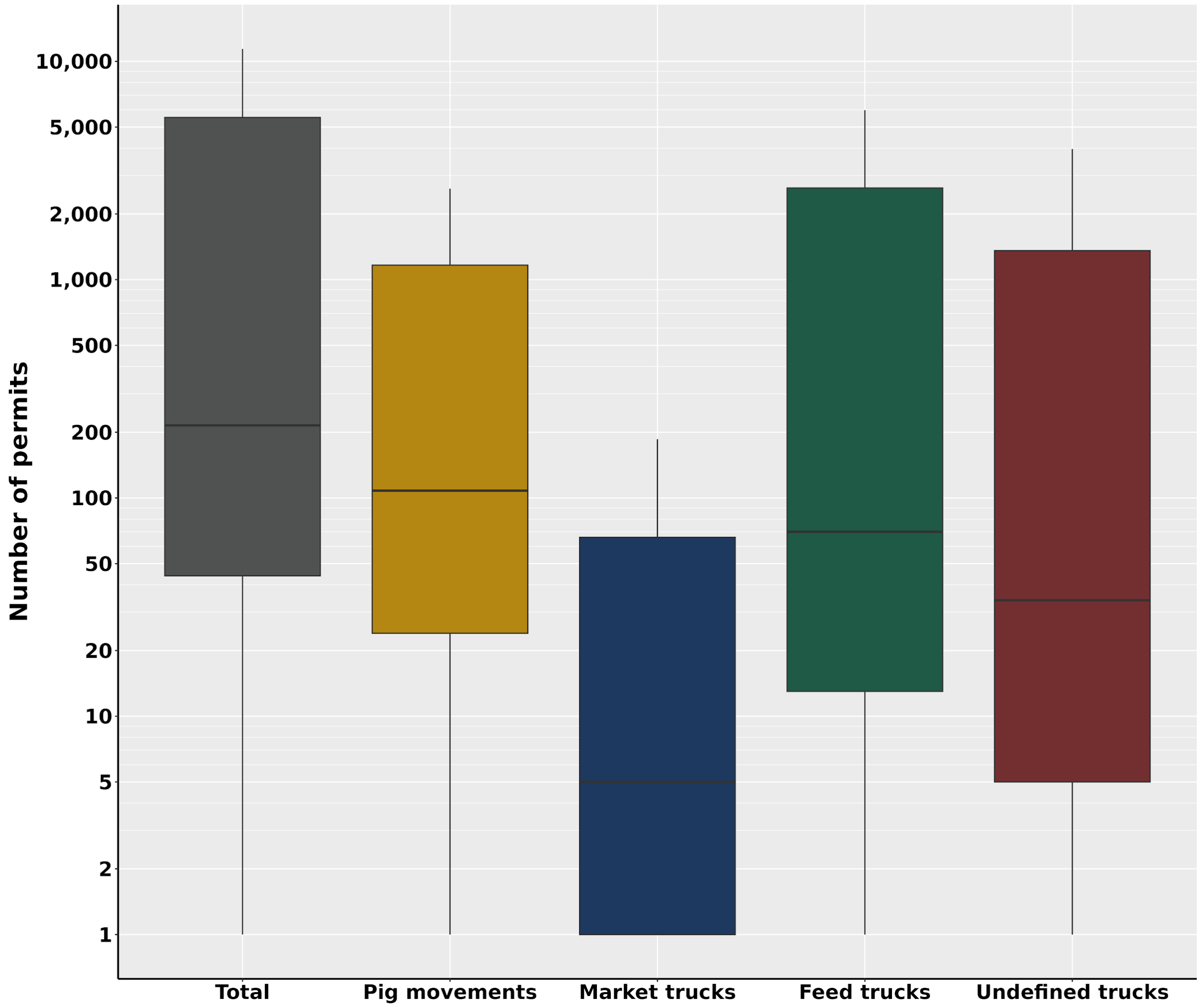


**Figure S16. Median cumulative permits required by 270 days of the epidemic, total and by movement type, under the NRP scenario.**

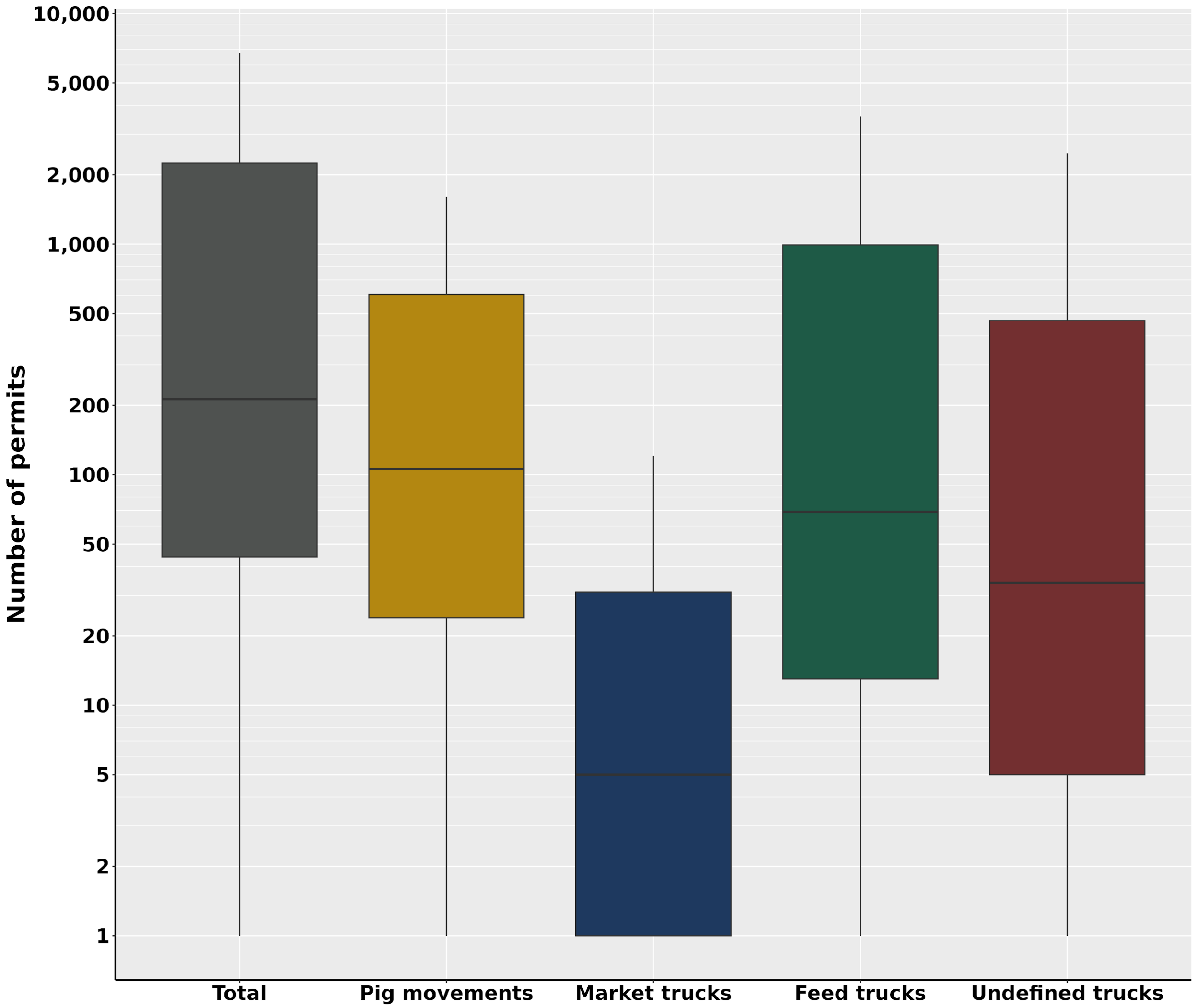


**Figure S17. Median cumulative permits required by 180 days of the epidemic, total and by movement type, under the NRP scenario.**

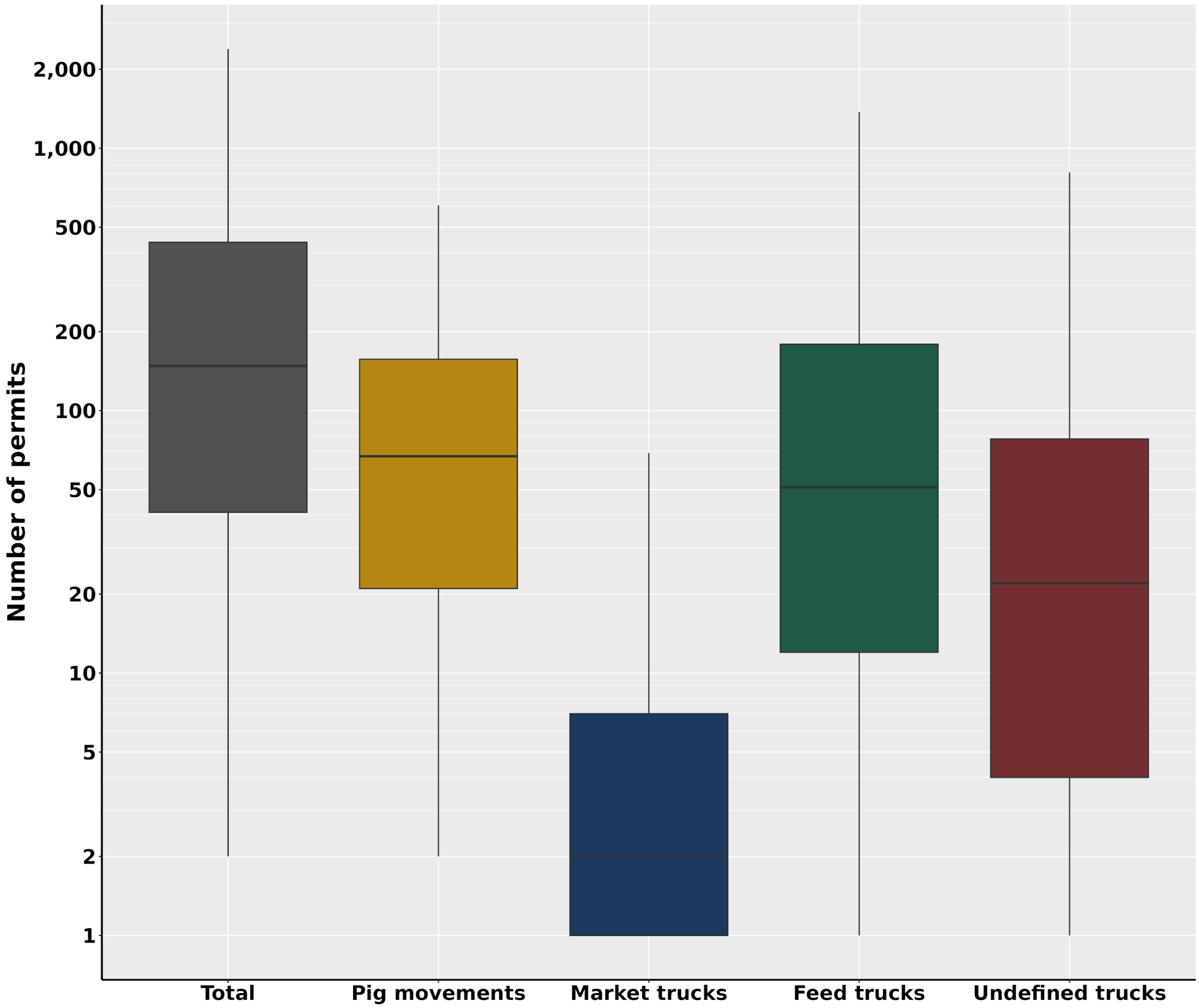


**Figure S18. Median cumulative permits required by 90 days of the epidemic, total and by movement type, under the NRP scenario.**

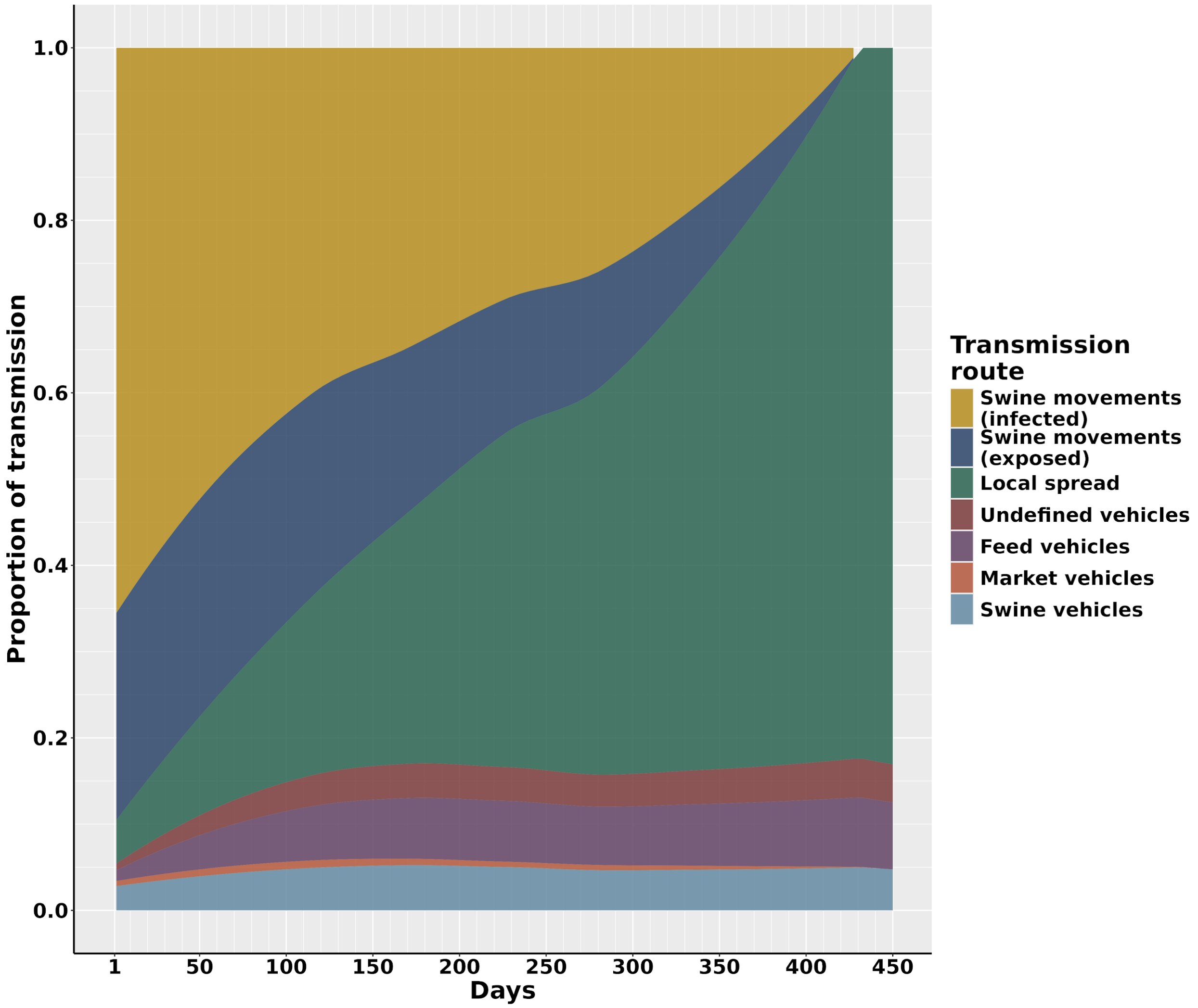


**Figure S19. Contribution of transmission routes to ASF infection over 450 days of the epidemic under the NRP scenario.** Day one of the epidemic is considered the day of introduction, transmission did not occur until day two.

Sample size calculations for surveillance testing

The sample sizes for each farm were calculated from information provided by the (USDA, 2020) and (Cannon, 2001). Equation 15 demonstrates the calculation.

$$sample\ size \approx\ a((1-(1-s)^{s(pq)^{-1}})(q-\frac{s(pq)-1}{2})) \qquad (15)$$

Where $s$ is the sensitivity of the diagnostic test, $p$ is the estimated prevalence of the disease, $q$ the population at the barn level and $a$ is the total number of barns on the farm.

**Table S4. Median (IQR) cumulative secondary infections by production type for the three, six, nine, and twelve-month elimination scenarios**

| *Production type* | *Three month scenario* | *Six month scenario* | *Nine month scenario* | *Twelve month scenario* | *NRP Scenario (Twelve Months)* |
|---|---|---|---|---|---|
| Sow | 0 (0 - 1) | 0 (0 - 1) | 0 (0 - 1) | 0 (0 - 1) | 1 (0 - 132) |
| Nursery | 0 (0 - 1) | 1 (0 - 2) | 1 (0 - 2) | 1 (0 - 2) | 2 (0 - 202) |
| Finisher | 1 (0 - 4) | 1 (0 - 5) | 1 (0 - 6) | 1 (0 - 6) | 4 (1 - 600) |
| Wean-to-finish | 0 (0 - 0) | 0 (0 - 1) | 0 (0 - 1) | 0 (0 - 1) | 1 (0 - 65) |
| Farrow-to-finish | 0 (0 - 0) | 0 (0 - 0) | 0 (0 - 0) | 0 (0 - 0) | 0 (0 - 0) |
| Boar stud | 0 (0 - 0) | 0 (0 - 0) | 0 (0 - 0) | 0 (0 - 0) | 0 (0 - 2) |
| Gilt | 0 (0 - 0) | 0 (0 - 0) | 0 (0 - 0) | 0 (0 - 0) | 0 (0 - 3) |
| Isolation | 0 (0 - 0) | 0 (0 - 0) | 0 (0 - 0) | 0 (0 - 0) | 0 (0 - 0) |

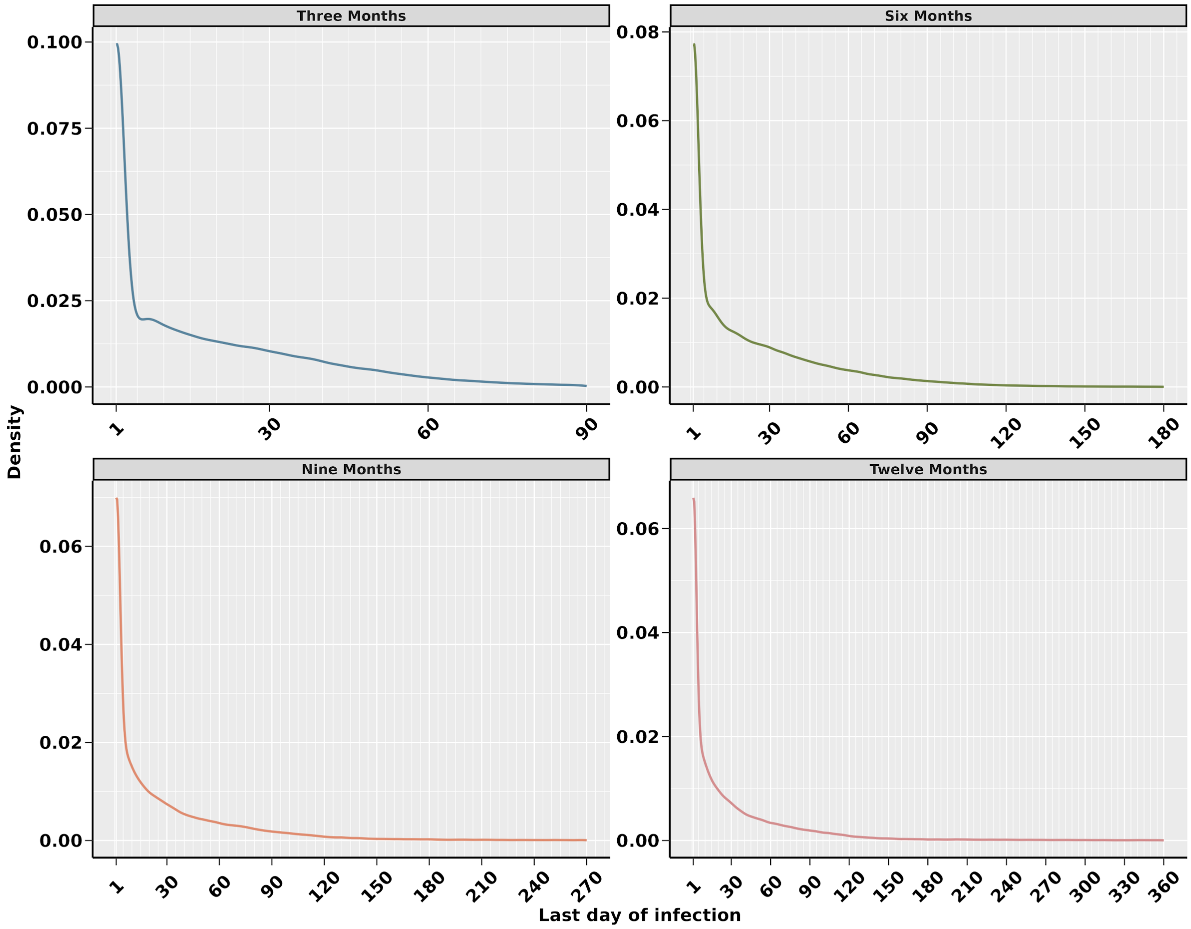


**Figure S20. Density plots showing the last day of infection for simulations in the three, six, nine and twelve month elimination scenarios.** We have only included the simulations that were controlled under the specified elimination time frame.

**Table S5. Percentage of simulations eliminated under each control strategy tested for twelve, nine, six and three months**

| *Control Strategy* | *Elimination % at twelve months* | *Elimination % at nine months* | *Elimination % at six months* | *Elimination % at three months* |
|---|---|---|---|---|
| **NRP Scenario** | **65.1** | **64.7** | **63.9** | **60.2** |
| Strategy #1 | 79.5 | 77.0 | 73.1 | 64.4 |
| Strategy #2 | 90.3 | 87.3 | 82.9 | 74.1 |
| Strategy #3 | 95.4 | 94.1 | 92.1 | 86.2 |
| Strategy #4 | 98.0 | 97.0 | 95.3 | 88.5 |
| **Twelve Month Scenario** (Strategy #5) | **99.1** | **98.4** | **97.0** | **90.5** |
| **Nine Month Scenario** (Strategy #6) | - | **99.0** | **97.9** | **91.8** |
| Strategy #7 | - | - | 98.6 | 92.1 |
| **Six Month Scenario** (Strategy #8) | - | - | **99.8** | **96.7** |
| Strategy #9 | - | - | - | 97.7 |
| Strategy #10 | - | - | - | 98.8 |
| **Three Month Scenario** (Strategy #11) | - | - | - | **99.2** |

"-" indicates that the strategy scenario was not ran for that time frame

**Table S6. Median (IQR) cumulative depopulated animals by production type for the three, six, nine and twelve month elimination scenarios**

| ***Production type*** | ***Three month scenario*** | ***Six month scenario*** | ***Nine month scenario*** | ***Twelve month scenario*** | ***NRP Scenario (Twelve Months)*** |
|---|---|---|---|---|---|
| Sow | 0 (0 - 2,600) | 0 (0 - 3,250) | 0 (0 - 3,600) | 0 (0 - 3,804) | 2,720 (0 - 337,730) |
| Nursery | 0 (0 - 6,400) | 2,328 (0 - 7,680) | 2,600 (0 - 8,240) | 2,600 (0 - 8,768) | 7,800 (0 - 836,852) |
| Finisher | 5,760 (0 - 18,080) | 5,880 (0 - 21,512) | 6,072 (0 - 25,440) | 6,120 (0 - 26,904) | 17,936 (0 - 2,268,341) |
| Wean-to-finish | 0 (0 - 0) | 0 (0 - 2,632) | 0 (0 - 4,100) | 0 (0 - 4,410) | 4,410 (0 - 389,284) |
| Farrow-to-finish | 0 (0 - 0) | 0 (0 - 0) | 0 (0 - 0) | 0 (0 - 0) | 0 (0 - 0) |
| Boar stud | 0 (0 - 0) | 0 (0 - 0) | 0 (0 - 0) | 0 (0 - 0) | 0 (0 - 283) |
| Gilt | 0 (0 - 0) | 0 (0 - 0) | 0 (0 - 0) | 0 (0 - 0) | 0 (0 - 2,000) |
| Isolation | 0 (0 - 0) | 0 (0 - 0) | 0 (0 - 0) | 0 (0 - 0) | 0 (0 - 0) |

**Table S7. Median (IQR) cumulative diagnostic tests by test reason for the three, six, nine and twelve month elimination scenarios**

| *Test reason* | *Three month scenario* | *Six month scenario* | *Nine month scenario* | *Twelve month scenario* | *NRP Scenario (Twelve Months)* |
|---|---|---|---|---|---|
| Direct contact | 588 (0 - 2,436) | 192 (0 - 888) | 212 (0 - 1,080) | 216 (0 - 1,152) | 288 (12 - 756) |
| Indirect contact | 5,684 (588 - 27,010) | 1,800 (192 - 10,558) | 1,896 (192 - 13,406) | 1,960 (192 - 14,348) | 1,981 (312 - 4,092) |
| Infected zone | 104,022 (38,158 - 189,127) | 31,785 (11,685 - 76,851) | 6,340 (2,160 - 18,802) | 2,621 (840 - 8,806) | 3,504 (660 - 66,198) |
| Buffer zone | 179,620 (97,608 - 224,342) | 69,248 (25,228 - 130,389) | 13,779 (4,860 - 35,182) | 10,830 (3,933 - 29,565) | 3,624 (744 - 21,547) |
| Surveillance zone | 31,730 (26,377 - 45,334) | 26,552 (14,450 - 39,712) | 6,951 (2,464 - 15,638) | 6,213 (2,175 - 15,149) | 4,470 (933 - 21,433) |

**Table S8. Median (IQR) cumulative permits by movement type for the three, six, nine and twelve month elimination scenarios**

| ***Movement type*** | ***Three month scenario*** | ***Six month scenario*** | ***Nine month scenario*** | ***Twelve month scenario*** | ***NRP Scenario (Twelve Months)*** |
|---|---|---|---|---|---|
| Pigs | 121 (0 - 554) | 834 (373 - 1,754) | 150 (62 - 493) | 109 (43 - 384) | 108 (24 - 1,878) |
| Market trucks | 133 (52 - 210) | 46 (11 - 101) | 9 (2 - 28) | 6 (1 - 22) | 5 (0 - 118) |
| Feed trucks | 1,731 (832 - 2,545) | 623 (193 - 1,240) | 136 (43 - 370) | 95 (30 - 274) | 70 (13 - 7,169) |
| Undefined trucks | 940 (479 - 1,447) | 338 (107 - 698) | 72 (21 - 203) | 50 (14 - 152) | 34 (5 - 4,253) |

**Table S9. Results of the LHS-PRCC sensitivity analysis including estimates, test statistic and p values**

| *Parameter* | *PRCC estimate* | *Lower bound* | *Upper bound* | *Test statistic* | *P value* |
|---|---|---|---|---|---|
| Transmission rate for infected pig movements | 0.0245 | -0.0032 | 0.0522 | 1.7339 | 0.0830 |
| Transmission rate for exposed pig movements | -0.0190 | -0.0467 | 0.0087 | -1.3442 | 0.1790 |
| Transmission rate for movements of pig trucks | 0.1105 | 0.0830 | 0.1381 | 7.8624 | < 0.001 |
| Transmission rate for movements of market trucks | 0.0447 | 0.0170 | 0.0724 | 3.1645 | < 0.01 |
| Transmission rate for feed delivery trucks | 0.3550 | 0.2390 | 0.3809 | 26.8430 | < 0.001 |
| Transmission rate for undefined trucks | 0.0810 | 0.0533 | 0.1086 | 5.7418 | < 0.001 |
| Maximum probability of transmission | -0.8217 | -0.8375 | -0.8059 | -101.9211 | < 0.001 |
| Gradient of transmission probability decline over distance | 0.6378 | 0.6165 | 0.6592 | 58.5511 | < 0.001 |
| Effective surveillance in sow farms | 0.0268 | -0.0009 | 0.0545 | 1.8934 | 0.0584 |
| Effective surveillance in nursery farms | 0.0063 | -0.0215 | 0.0340 | 0.4430 | 0.6578 |
| Effective surveillance in finisher farms | 0.0304 | 0.0027 | 0.0582 | 2.1532 | < 0.05 |
| Local transmission spread cut off | 0.5635 | 0.5406 | 0.5864 | 48.2236 | < 0.001 |
| Latent period of virus | -0.1630 | -0.1904 | -0.1357 | -11.6830 | < 0.001 |
| Time to detection midpoint | -0.0095 | -0.0373 | 0.0182 | -0.6742 | 0.5002 |

Calculated with 4998 degrees of freedom. Values have be rounded to four decimal points

**Table S10. Median number of farms within the radius of the control area and surveillance zone at different sizes**

| *Total distance of control area and surveillance zone combined (km)* | *Median number of farms (IQR)* |
|---|---|
| 10 | 46 (14 - 72) |
| 13 | 71 (22 115) |
| 15 | 92 (27 - 150) |
| 30 | 331 (108 - 488) |
| 45 | 675 (233 - 867) |
| 60 | 1,008 (498 - 1,227) |

USDA, 2020. African swine fever response plan: The Red Book.
